# EstG is a novel esterase required for cell envelope integrity

**DOI:** 10.1101/2022.04.12.488081

**Authors:** Allison K. Daitch, Benjamin C. Orsburn, Zan Chen, Laura Alvarez, Colten D. Eberhard, Kousik Sundararajan, Rilee Zeinert, Dale F Kreitler, Jean Jakoncic, Peter Chien, Felipe Cava, Sandra B. Gabelli, Namandjé N. Bumpus, Erin D. Goley

## Abstract

Proper regulation of the bacterial cell envelope is critical for cell survival. Identification and characterization of enzymes that maintain cell envelope homeostasis is crucial, as they can be targets for effective antibiotics. In this study, we have identified a novel enzyme, called EstG, whose activity protects cells from a variety of lethal assaults in the ⍺-proteobacterium *Caulobacter crescentus*. Despite homology to transpeptidase family cell wall enzymes and an ability to protect against cell wall-targeting antibiotics, EstG does not demonstrate biochemical activity towards cell wall substrates. Instead, EstG is genetically connected to the periplasmic enzymes OpgH and BglX, responsible for synthesis and hydrolysis of osmoregulated periplasmic glucans (OPGs), respectively. The crystal structure of EstG revealed similarities to esterases and transesterases, and we demonstrated esterase activity of EstG *in vitro*. Using biochemical fractionation, we identified a cyclic hexamer of glucose as a likely substrate of EstG. This molecule is the first OPG described in *Caulobacter* and establishes a novel class of OPGs, the regulation and modification of which is important for stress survival and adaptation to fluctuating environments. Our data indicate that EstG, BglX, and OpgH comprise a previously unknown OPG pathway in *Caulobacter*. Ultimately, we propose that EstG is a novel enzyme that, instead of acting on the cell wall, acts on cyclic OPGs to provide resistance to a variety of cellular stresses.

## Introduction

The bacterial cell envelope is a multi-component structure that protects bacteria from the external environment. The envelope is an essential physical barrier to the surroundings, and the factors responsible for building and maintaining the envelope are therefore ideal targets for antibiotics. The Gram-negative cell envelope consists of the inner and outer membranes, with the periplasmic space (or periplasm) between them (Silhavy et al., 2010). The bacterial cell wall, made of peptidoglycan (PG), forms a protective meshwork in the periplasm that prevents cell lysis due to turgor pressure (Huang et al., 2008). During growth and division, essential PG metabolic enzymes synthesize, modify, and hydrolyze the PG. Two major classes of PG synthetic enzymes include the glycosyltransferases and transpeptidases (TPases), which catalyze polymerization of the glycan strands and crosslinking of strands via the peptide stems, respectively (Daitch and Goley, 2020). For almost all bacteria, PG is an essential structural component, and the primary biosynthetic PG enzymes are essential during normal growth and division (Daitch and Goley, 2020). Because of this function, these enzymes are the targets of bactericidal antibiotics, such as β-lactams, which inhibit the TPase activity of penicillin-binding proteins (PBPs) (Fisher and Mobashery, 2020). Though some of the most effective antibiotic targets are PG enzymes, disruption of other components of the envelope can also sensitize cells to stress or antibiotics (May and Grabowicz, 2018; Sutterlin et al., 2016). Thus, understanding the elements of the cell envelope and their relationships to each other is crucial for identifying new drug targets.

In addition to PG, the periplasm of proteobacteria may contain glycopolymers important for maintaining cell envelope integrity called osmoregulated periplasmic glucans (OPGs, also called membrane-derived oligosaccharides). OPGs are glucose polymers that are made in the periplasm and are thought to function as osmoprotectants in response to changes in the environment (Bontemps-Gallo et al., 2017). Across Gram-negative species, OPGs vary in size, ranging from 5 to 24 glucose units, and geometry, exhibiting linear, branched, and/or cyclic structures depending on which OPG metabolic enzymes are encoded in a given organism (Bohin, 2000; Bontemps-Gallo et al., 2017). OPGs may also be modified with, for example, phospholipid moieties (e.g. phosphoglycerol) or products of intermediary metabolism (e.g. succinyl), which can influence the polymer’s overall charge (Bohin, 2000; Bontemps-Gallo et al., 2017). Previously characterized OPGs in α-proteobacteria are large (10-25 glucose units), cyclic, and highly modified (Bohin, 2000). In some bacteria, OPGs are implicated in stress tolerance, as disruption of OPG genes results in increased sensitivity to antibiotics and cell envelope stresses (Bontemps-Gallo et al., 2017; Bontemps-Gallo and Lacroix, 2015). Despite decades of research on OPGs, we have limited knowledge about the diversity of OPG structures, modifications, and metabolic enzymes across bacteria, suggesting the possibility of undiscovered OPG molecules, pathways, and functions.

In this study, we sought to identify factors required to survive cell wall stress in the α-proteobacterium *Caulobacter crescentus*, which is a well-studied model for morphogenesis and PG metabolism (Woldemeskel and Goley, 2017). We used a genetic screen to identify an uncharacterized protein required for survival during cell wall stress that we called EstG (*E*sterase for *S*tress *T*olerance acting on *G*lucans, described below). Although EstG is annotated as a member of the TPase superfamily, which consists primarily of PG-acting enzymes, it has no detectable activity towards PG. Our data indicate that EstG instead acts in the OPG metabolic pathway in *Caulobacter,* as it has genetic interactions with the putative OPG enzymes BglX, a periplasmic glucohydrolase, and OpgH, the OPG synthase. The crystal structure of EstG revealed similarity to esterases and transesterases and we confirmed esterase activity *in vitro.* An unbiased mass spectrometry approach identified a native substrate of EstG as a periplasmic, cyclic hexamer of glucose. This is the first OPG identified in *Caulobacter* and establishes a new class of OPGs in α-proteobacteria. We propose that EstG is a novel enzyme that, instead of acting on the PG like most other well-characterized members of the TPase superfamily, acts on cyclic OPGs to fortify the cell envelope and provide resistance to a variety of cellular stresses.

## Results

### EstG is essential for suppression of toxic cell wall misregulation

This study initiated with our interest in understanding *Caulobacter* PG metabolism during cell division, which is orchestrated by the polymerizing tubulin homolog FtsZ. We previously demonstrated that expression of a mutant of *ftsZ* lacking the C-terminal linker domain (called ΔCTL) results in misregulation of PG enzymes and cell death, similar to the effects of β-lactam antibiotic treatment (Figure 1A) (Sundararajan et al., 2015). We leveraged ΔCTL toxicity to understand mechanisms of stress survival. To this end, we conducted a screen to identify spontaneous suppressors of ΔCTL-induced lethality (Figure 1A) (Woldemeskel et al., 2020). Whole genome sequencing of suppressors revealed mutations in genes largely involved in nutrient stress responses (i.e., *spoT* (Boutte and Crosson, 2011), *cdnL* (Woldemeskel et al., 2020; Gallego-Garcia et al., 2017), and *phoB* (Lubin et al., 2016)) (Figure 1A, Supplemental Table 1). Although each of the ΔCTL suppressors reduce growth rate on their own, slow growth was not sufficient to suppress ΔCTL-induced lethality as tested by growth at low temperature, or in the presence of sub-lethal doses of chloramphenicol or fosfomycin to reduce translation or PG synthesis respectively (Figure 1—Figure supplement 1A-F). This indicated that these mutations suppressed ΔCTL through other mechanisms. We were especially intrigued by the identification of suppressing mutations in *spoT*, since SpoT is the primary mediator of the stringent response in *Caulobacter* and the stringent response has been implicated in antibiotic resistance (Boutte and Crosson, 2011).

**Figure 1:**
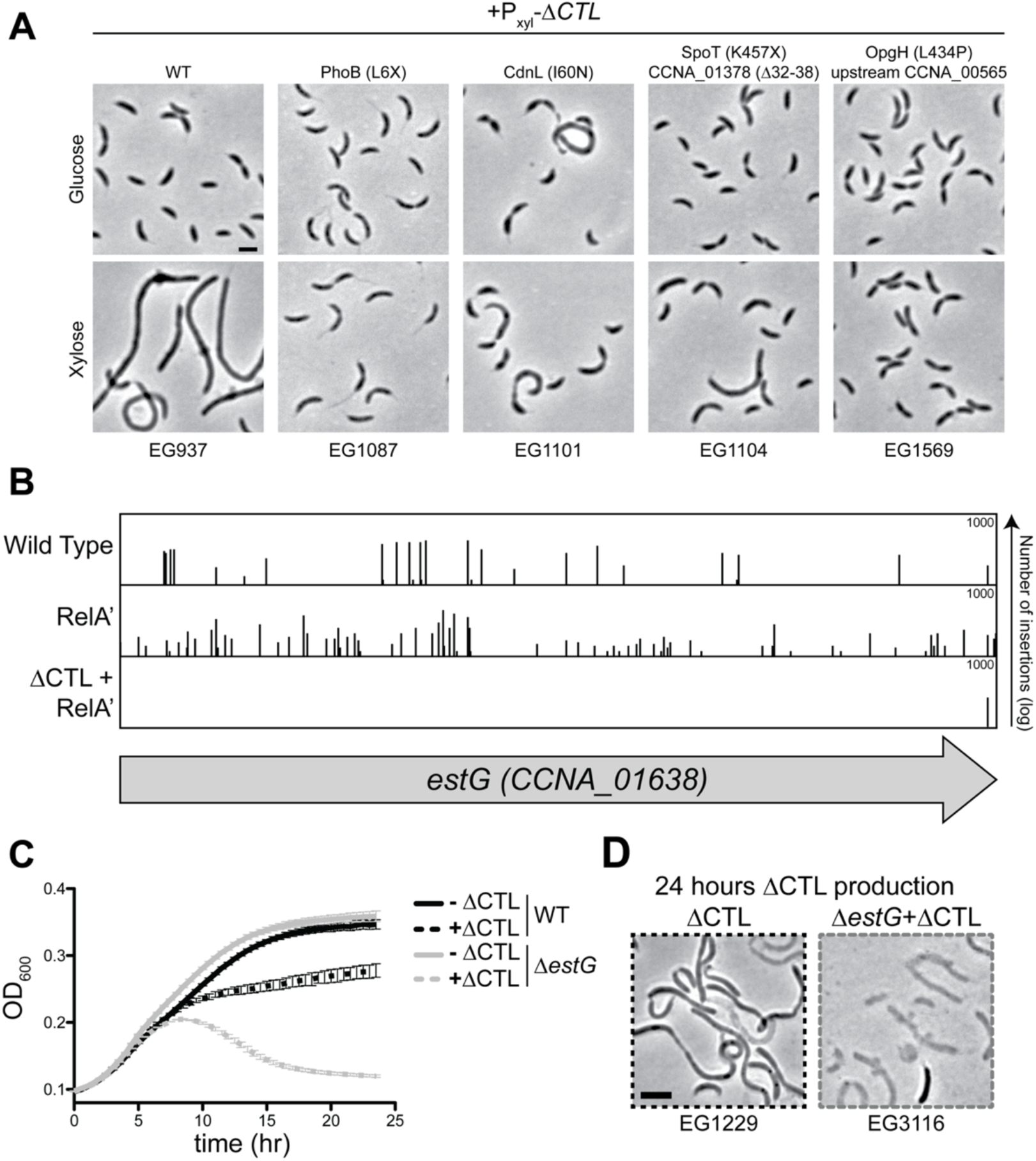
EstG is required to suppress ΔCTL-mediated lethality. **A.** Phase contrast images of ΔCTL and suppressors +/- ΔCTL production. Indicated strains are grown with 0.3% glucose (-ΔCTL) or 0.3% xylose (+ΔCTL) for 7 hours before imaging. Scale bar, 2 µm. Amino acid X represents a premature stop codon. **B.** Line plot of transposon insertion frequency along the gene locus for *CCNA_01638* (named *estG*) as determined by transposon sequencing (Tn-Seq) analysis in wild type (WT; EG865), high (p)ppGpp production (RelA’, EG1799), and ΔCTL with high (p)ppGpp production (ΔCTL+RelA’, EG1616). **C.** Growth curve of strains EG1229 (WT) and EG3116 (Δ*estG)* with and without ΔCTL production (+/- 0.3% xylose) as monitored by OD_600_. **D.** Phase contrast images of WT and Δ*estG* from the 24-hour timepoint of the growth curve in panel C. Scale bar, 2 µm.

We determined that SpoT-mediated suppression of ΔCTL was a result of high levels of the signaling alarmone (p)ppGpp using an inducible and constitutively activated form of the *Escherichia coli* (*E. coli*) (p)ppGpp synthase, RelA (hereafter called RelA’) (Figure 1—Figure supplement 1G) (Gonzalez and Collier, 2014). To better understand how high levels of (p)ppGpp suppress ΔCTL-induced lethality, we conducted comparative transposon sequencing (Tn-Seq) to identify genes that were synthetically lethal with ΔCTL expression, using the following strains: wild type (WT), RelA’-producing, and RelA’-producing with ΔCTL. Notably, we identified a gene, *CCNA_01638* (hereafter named *estG* for *E*sterase for *S*tress *T*olerance acting on *G*lucans, for reasons described below), that appeared to be essential only in the presence of ΔCTL stress (Figure 1B). *estG* acquired abundant transposon insertions in WT and RelA’ backgrounds, suggesting that it is non-essential in those strains. However, there were almost no transposon insertions in *estG* in RelA’-producing cells that also produced ΔCTL, indicating an essential function of EstG in the presence of ΔCTL (Figure 1B). EstG is an uncharacterized protein that is annotated as a β-lactamase family protein in the transpeptidase superfamily, which primarily consists of PG enzymes. We were therefore interested in studying EstG and its relationship to surviving PG stress.

To validate our Tn-Seq findings, we deleted *estG* in a strain with xylose-inducible production of ΔCTL (Figure 1C-D). This strain grew comparably to a ΔCTL uninduced strain in the absence of xylose (Figure 1C, solid lines). Production of ΔCTL in an otherwise WT background resulted in cell filamentation and lysis over time, as expected (Figure 1C-D, black). Notably, producing ΔCTL in a Δ*estG* background resulted in faster cell lysis compared to ΔCTL in a WT background (Figure 1C-D, grey). We were struck by the importance of *estG* in the presence of ΔCTL-induced stress and sought to further understand the function of EstG.

### *estG* is non-essential in unstressed conditions, but required for survival during cell wall stress

Our Tn-Seq results, as well as prior Tn-Seq data, indicated that *estG* would be non-essential in an otherwise WT background (Figure 1B) (Christen et al., 2011). We confirmed this by generating a deletion of *estG* (Δ*estG*) and comparing its growth and morphology to WT. We confirmed deletion of *estG* via western blotting with an affinity-purified EstG antibody (Figure 2—Figure supplement 1A). Δ*estG* cells grew comparably to WT by optical density (Figure 2A) and spot dilution (Figure 2B), though the colony size of the Δ*estG* strain is slightly smaller than WT. Additionally, by phase contrast microscopy, Δ*estG* cells look morphologically identical to WT (Figure 2C). Therefore, *estG* is non-essential under normal growth conditions, but becomes essential during the cell wall stress induced by ΔCTL.

**Figure 2:**
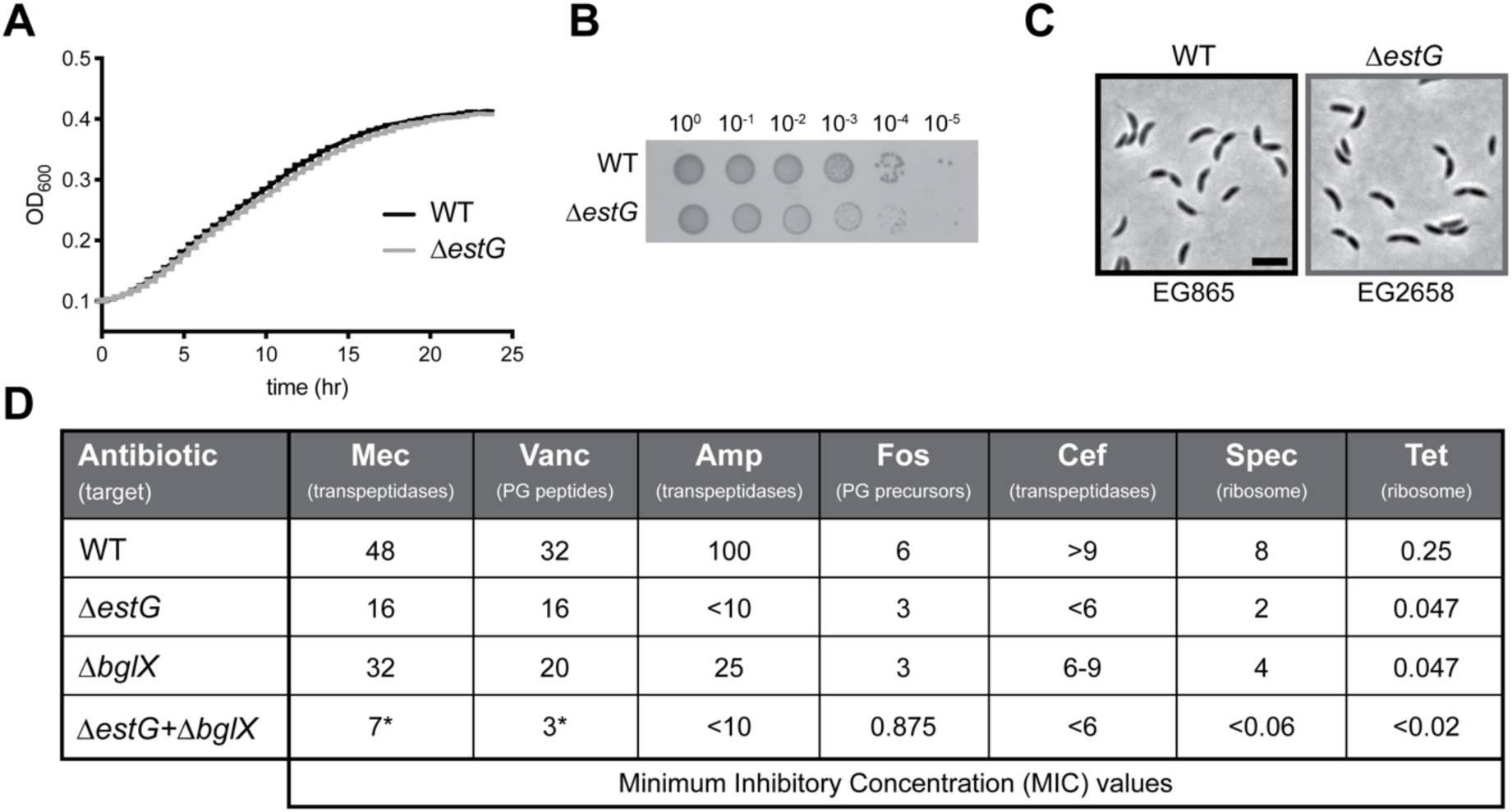
Δ*estG* does not impact cell viability or growth in unstressed conditions. **A.** Growth curve, **B.** spot dilutions, and **C.** phase contrast images of wild type (WT, EG865) and Δ*estG* (EG2658). Culture dilutions are as indicated. Scale bar, 2 µm. **D.** Minimum inhibitory concentrations (MIC) of WT (EG865), *ΔestG* (EG2658), Δ*bglX* (EG3279), and Δ*estGΔbglX* (EG3282) against peptidoglycan (PG)- and ribosome-targeting antibiotics. Measurements in µg/mL. Mec=mecillinam; Vanc=vancomycin; Amp=ampicillin; Fos=fosfomycin; Cef=cephalexin; Spec=spectinomycin; Tet=tetracycline. Asterisk (*) represents value with a secondary zone of light inhibition.

After observing the essentiality of *estG* during ΔCTL production, we hypothesized that EstG may also be required to survive other cell wall stresses, such as cell wall-targeting antibiotics. To test this, we measured the minimum inhibitory concentrations (MIC) of a variety of antibiotics against WT and Δ*estG* cells (Figure 2D). Δ*estG* was hypersensitive to every cell wall antibiotic tested (mecillinam, vancomycin, ampicillin, fosfomycin, and cephalexin) compared to WT, represented by a decreased MIC value. To confirm that hypersensitivity was specifically attributable to loss of EstG, we complemented with a vanillate-inducible copy of *estG* and showed that resistance to ampicillin was restored (Figure 2—Figure supplement 1B). This indicates a broadly important role of EstG during cell wall stress.

While exploring the possible role of EstG, we noticed the gene immediately downstream from *estG*, *CCNA_01639*, is also annotated as a β-lactamase family protein and we wondered if the two might be functionally related. CCNA_01639 has high sequence identity to EstG (52%), and both are predicted to reside in the periplasm (Juan et al., 2019). Despite similarity to EstG, however, deletion of *CCNA_01639* did not result in hypersensitivity to the β-lactam antibiotics ampicillin or cephalexin (Figure 2—Figure supplement 1C). Moreover, the double deletion, Δ*estGΔCCNA_01639*, phenocopied the single Δ*estG* mutant (Figure 2—Figure supplement 1C). Since Δ*CCNA_01639* had no detectable phenotype or genetic relationship to *estG*, we focused the remainder of our study on characterizing EstG.

### EstG is periplasmic with no detectable cell wall activity

EstG is 462 amino acids and has an N-terminal putative signal sequence, with cleavage predicted between residues 30 and 31 (Juan et al., 2019). To study the periplasmic localization of EstG, we expressed an inducible EstG*-*β-lactamase (EstG-BlaM) fusion protein in an otherwise β-lactamase deficient strain (Δ*blaA*; BlaA is the primary β-lactamase that confers β-lactam resistance to *Caulobacter* (West et al., 2002)). These cells will only be resistant to ampicillin if EstG contains a periplasmic signal sequence to transport the fused β-lactamase to the periplasm (Möll et al., 2010). The EstG-BlaM strain, when plated in the presence of inducer, displayed resistance to ampicillin, thus validating the predicted periplasmic localization of EstG (Figure 2— Figure Supplement 1D).

The classification of EstG as a β-lactamase family protein as well as the hypersensitivity of Δ*estG* to ΔCTL and PG-targeting antibiotics suggested that EstG might act as a β-lactamase. However, purified EstG displayed negligible activity against nitrocefin, a substrate used to detect β-lactamase activity *in vitro* (Figure 2—Figure supplement 1E), compared to a *Caulobacter* enzyme with moderate β-lactamase activity, EstA (Ryu et al., 2016). This, however, does not rule out an activity against the cell wall, so we next tested for ability to bind to the cell wall. *In vitro,* purified EstG pelleted with PG isolated from WT *Caulobacter*, whereas a non-cell wall binding protein (glutathione S-transferase, GST) remained soluble (Figure 2—Figure supplement 1F). This demonstrates the ability of EstG to bind to some component of the PG. Despite this, EstG did not have detectable activity against any of the most abundant muropeptide species (M4, M5, D44, and D45) or purified PG sacculi *in vitro* (Figure 2—Figure supplement 2A-G). Finally, we asked if we could identify EstG-dependent chemical changes in PG via muropeptide analysis of sacculi isolated from Δ*estG* cells as compared to WT. Again, there were no significant differences between Δ*estG* and WT PG (Figure 2—Figure supplement 2H-I, Table 1). This was surprising given the classification of EstG as a transpeptidase superfamily enzyme, consisting of TPases and carboxypeptidases, which often have detectable activity on cell wall substrates. Considering EstG’s lack of activity against cell wall substrates *in vitro*, we hypothesized that EstG’s substrate is novel and not directly related to PG metabolism.

**Table 1.** Deletion of *estG* does not alter the muropeptide profile of *Caulobacter*. Table outlines muropeptide relative molar abundance (%). GlcNAc: N-Acetyl glucosamine. MurNAc: N-Acetyl muramic acid. Ala: Alanine. Glu: Glutamic acid. mDAP: meso-diaminopimelic acid. Gly: Glycine. Statistical analysis performed using t-test analysis. * = P < 0.05 and > 10% variation compared to WT.

| Peak | Muropeptide | Structure | WT | $\Delta estG$ |
| --- | --- | --- | --- | --- |
| 1 | M3 | GlcNAc-MurNAc-L-Ala-D-Glu-mDAP | 0.42 | 0.3 |
| 2 | M4<sup>G</sup> | GlcNAc-MurNAc-L-Ala-D-Glu-mDAP-Gly | 0.37 | 0.28 |
| 3 | M5<sup>G</sup> | GlcNAc-MurNAc-L-Ala-D-Glu-mDAP-D-Ala-Gly | 10.32 | 10.17 |
| 4 | M4 | GlcNAc-MurNAc-L-Ala-D-Glu-mDAP-D-Ala | 30.01 | 29.48 |
| 5 | M2 | GlcNAc-MurNAc-L-Ala-D-Glu | 1.16 | 0.71 |
| 6 | M5 | GlcNAc-MurNAc-L-Ala-D-Glu-mDAP-D-Ala-D-Ala | 19.08 | 18.26 |
| 7 | D45<sup>G</sup> | M4-M5G (DD-crosslink) | 4.69 | 4.75 |
| 8 | D44 | M4-M4 (DD-crosslink) | 7.79 | 7.78 |
| 9 | M5<sup>G</sup> Anh | GlcNAc-(1-6anhydro)MurNAc-L-Ala-D-Glu-mDAP-D-Ala-Gly | 0.71 | 0.69 |
| 10 | D45 | M4-M5 (DD-crosslink) | 7.02 | 6.98 |
| 11 | M4<sup>Anh</sup> | GlcNAc-(1-6anhydro)MurNAc-L-Ala-D-Glu-mDAP-D-Ala | 0.75 | 0.72 |
| 12 | T445<sup>G</sup> | M4-M4-M5<sup>G</sup> (DD-crosslink) | 0.63 | 0.79 |
| 13 | T444 | M4-M4-M4 (DD-crosslink) | 1.1 | 1.07 |
| 14 | T445 | M4-M4-M5 (DD-crosslink) | 0.87 | 0.85 |
| 15 | D45<sup>G</sup> Anh | M4-M5<sup>G</sup> Anh (DD-crosslink) | 0.73 | 0.71 |
| 16 | D44<sup>Anh</sup> | M4-M4<sup>Anh</sup> (DD-crosslink) | 2.32 | 2.94 |
| 17 | D45<sup>Anh</sup> | M4-M5<sup>Anh</sup> (DD-crosslink) | 2.04 | 2.69 |
| 18 | T445<sup>G</sup> Anh | M4-M4-M5<sup>G</sup> Anh (DD-crosslink) | 1.68 | 1.81 |
| 19 | T444<sup>Anh</sup> | M4-M4-M4<sup>Anh</sup> (DD-crosslink) | 2.72 | 2.97 |
| 20 | T445<sup>Anh</sup> | M4-M4-M5<sup>Anh</sup> (DD-crosslink) | 1.88 | 2.04 |
| 21 | T445<sup>G</sup> Anh,Anh | M4-M4<sup>Anh</sup>-M5<sup>G</sup> Anh (DD-crosslink) | 0.54 | 0.69 |
| 22 | T444<sup>Anh,Anh</sup> | M4-M4<sup>Anh</sup>-M4<sup>Anh</sup> (DD-crosslink) | 2.46 | 2.44 |
| 23 | T445<sup>Anh,Anh</sup> | M4-M4<sup>Anh</sup>-M5<sup>Anh</sup> (DD-crosslink) | 0.73 | 0.9 |

### *estG* interacts genetically with *opgH*, which encodes a putative OPG synthase

To search for the molecular function of EstG in an unbiased fashion, we isolated and characterized spontaneous suppressors of the ampicillin sensitivity of Δ*estG*. We sequenced four suppressors total (Supplemental Table 1), but were most intrigued by a suppressing mutation in the essential gene, *opgH*, a periplasmic glucan glucosyltransferase (OpgH_L480P_) (Figure 3A). OpgH has been characterized in other organisms as the synthase of osmoregulated periplasmic glucans (OPGs) (Bontemps-Gallo et al., 2017). By BLAST searching, OpgH is the only homolog of known OPG-biosynthetic enzymes encoded in the *Caulobacter* genome, but the presence of OpgH indicates the existence of an undiscovered OPG pathway. Isolation of a suppressing mutation in OpgH led us to hypothesize that the sensitivities of *ΔestG* could be related to OPG production or modification.

**Figure 3:**
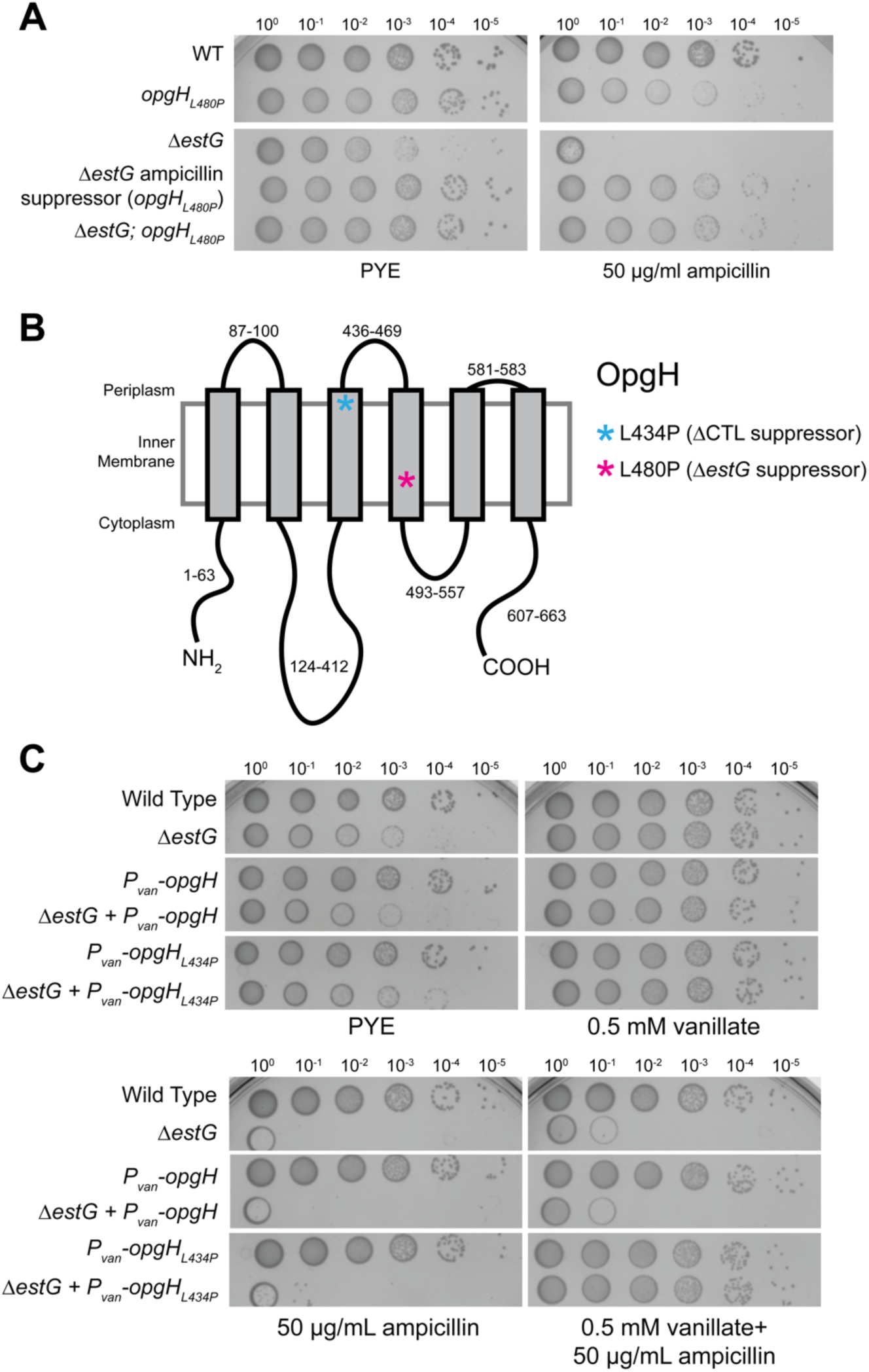
*opgH_L480P_* and *opgH_L434P_* suppress Δ*estG* sensitivities. **A.** Spot dilutions of WT (EG865), *opgH_L480P_* (EG3369), Δ*estG* (EG2658), *ΔestG* ampicillin suppressor (EG3105), *ΔestG*; *opgH_L480P_* (EG3371) grown on PYE agar alone or with 50 µg/mL ampicillin. Culture dilutions are as indicated. **B.** Schematic diagraming predicted topology of OpgH with grey boxes representing transmembrane domains and corresponding amino acids labeled. Asterisks represent approximate location of suppressing point mutations from the ΔCTL (EG1569) and *ΔestG* suppressors (EG3105). **C.** Spot dilutions of indicated strains on PYE agar alone or with added 0.5 mM vanillate and/or 50 µg/mL ampicillin. Strains are WT (EG865), Δ*estG* (EG2658), *P_van_-opgH* (EG3375), *ΔestG* + *P_van_-opgH* (EG3377), *P_van_-opgH_L434P_* (EG3577), and *ΔestG* + *P_van_-opgH_L434P_* (EG3579).

To characterize the suppressing mutation in *opgH*, we generated the suppressing mutation (*opgH_L480P_*) in a clean genetic background, in the presence or absence of *estG*. In the absence of stress, *opgH_L480P_* did not impact growth, but did restore Δ*estG* cells to a WT colony size (Figure 3A). In the presence of ampicillin, *opgH_L480P_* completely restored growth in a Δ*estG* background (Figure 3A). We also note that the *opgH_L480P_* mutation in a WT background exhibited moderate growth defects in the presence of ampicillin.

We hypothesized that the *opgH_L480P_* mutation might result in a loss of function variant, as the proline substitution is located within a predicted transmembrane domain (Figure 3B) (Krogh et al., 2001; Sonnhammer and Krogh, 2008). To ensure that the L480P mutation was not destabilizing the protein, we assessed the steady state levels of a 3x-Flag tagged version of the L480P mutant expressed from the native *opgH* locus and saw no difference in protein levels compared to WT (Figure 3—Figure supplement 1A). We then tested if OpgH_L480P_ could suppress Δ*estG* sensitivity to stress in the presence of WT OpgH by expressing vanillate-inducible *opgH_L480P_*. Indeed, expression of *opgH_L480P_* suppressed Δ*estG* sensitivity to ampicillin in a dominant fashion (Figure 3—Figure supplement 1B). These data led us to conclude that the OpgH_L480P_ mutant suppresses Δ*estG* by altering OpgH activity or function and is not a loss of function variant.

Interestingly, in revisiting our original ΔCTL suppressors, we discovered an independent suppressing mutation in *opgH* that restored growth in the presence of ΔCTL (Supplemental Table 1). This mutant, OpgH_L434P_ (Figure 1A), is also a leucine to proline mutation and is located at the edge of a different predicted transmembrane domain (Figure 3B). We tested if, like L480P, the L434P mutant could suppress Δ*estG* sensitivity in a dominant fashion. Strikingly, the OpgH_L434P_ mutant completely restored growth of Δ*estG* in the presence of ampicillin (Figure 3C). Collectively, our suppressor analyses solidify a genetic link between *estG* and *opgH*.

### *estG* and *bglX* are synthetically sick

To gain further insight into which pathway(s) EstG may impact, we examined *estG* on the Fitness Browser database (Wetmore et al., 2015). This database includes sensitivities of a genome-wide library of transposon mutants in *Caulobacter* to numerous stress and environmental conditions and reports on each gene’s mutant fitness profile. This resource reflected Δ*estG*’s sensitivities to cell wall antibiotics and also revealed genes that share a similar sensitivity profile to Δ*estG* when disrupted (i.e., genes that are “co-fit”). The top hit for co-fitness with *estG* was an uncharacterized gene, *bglX (CCNA_01162)*, predicted to encode a β-D-glucoside glucohydrolase. The BglX homolog in *Pseudomonas aeruginosa* (*P. aeruginosa*) cleaves glucose polymers (including OPGs) *in vitro*, but BglX homologs are otherwise uncharacterized, with little known about their physiological functions (Mahasenan et al., 2020).

We tested for activity of purified *Caulobacter* BglX as a glucohydrolase *in vitro* against the reporter substrate 4-nitrophenyl-β-D-glucopyranoside (pNPG), where hydrolysis of pNPG results in a color change that can be measured as absorbance over time (Mahasenan et al., 2020). BglX was able to hydrolyze pNPG in a concentration dependent manner, confirming its activity as a glucohydrolase (Figure 4A), while EstG displayed no activity against pNPG (Figure 4—Figure supplement 1A). *In vivo,* we determined that *bglX* is non-essential and that its loss does not appreciably affect growth or morphology (Figure 4B-C). As predicted, however, we found that Δ*bglX* shares all of the antibiotic sensitivities we observed for Δ*estG* (Figure 2D). We also confirmed periplasmic localization of BglX (Figure 4—Figure supplement 1B). Their similar sensitivity profiles indicated a possible genetic interaction between *estG* and *bglX*. Indeed, when both *estG* and *bglX* were deleted (Δ*estG*Δ*bglX*), cells had a growth defect and exhibited slight cell filamentation in unstressed conditions when compared to WT or either single deletion mutant (Figure 4B-C). The double deletion also had a lower MIC for all tested antibiotics compared to either of the single deletions, confirming a synthetic sickness between *estG* and *bglX* (Figure 2D).

**Figure 4:**
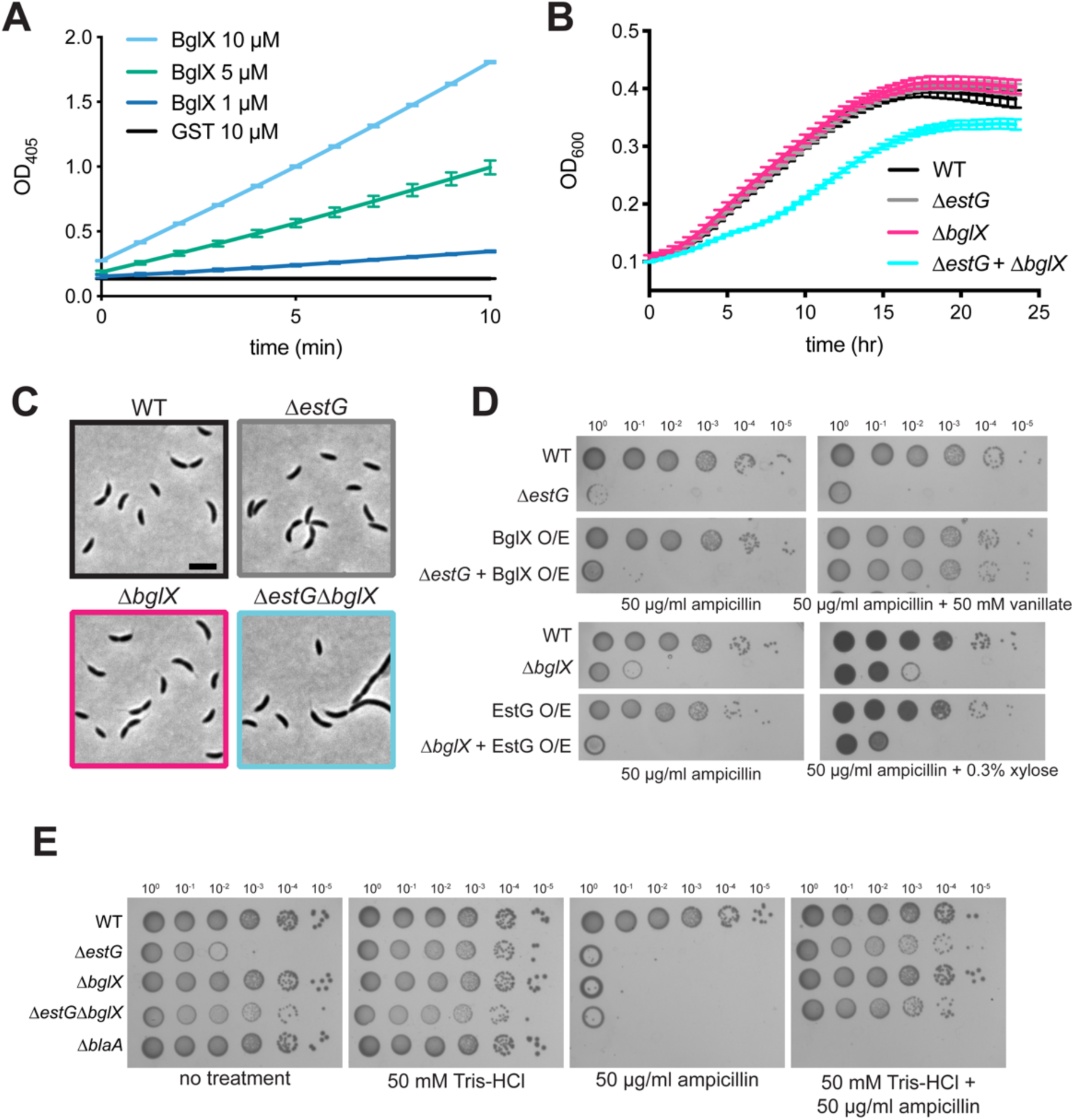
BglX is a glucosidase that interacts genetically with *estG*. **A.** 4-Nitrophenyl-β-D-glucopyranoside (pNPG) hydrolysis assay with purified BglX or GST at indicated amounts measured at OD_405_. **B.** Growth curve and **C.** phase contrast images of WT (EG865), Δ*estG* (EG2658), *ΔbglX* (EG3279), and *ΔestGΔbglX* (EG3282). Scale bar, 2 µm. **D.** Spot dilutions on PYE agar with 50 µg/mL ampicillin +/- 50 mM vanillate or 0.3% xylose of WT (EG865), Δ*estG* (EG2658), *P_van_-bglX* (BglX O/E, EG3384), Δ*estG+ P_van_-bglX* (EG3385), Δ*bglX* (EG3279), *P_xyl_-estG* (EG2759), and Δ*bglX+ P_xyl_-estG* (EG3425). **E.** Spot dilutions on PYE agar alone (no treatment) or with added 50 mM Tris-HCl and/or 50 µg/mL ampicillin of WT (EG865), Δ*estG* (EG2658), *ΔbglX* (EG3279), *ΔestGΔbglX* (EG3282), and Δ*blaA* (EG2408). Culture dilutions are as indicated.

From this synthetic interaction, we hypothesized that EstG and BglX fulfill a similar function. If so, overexpression of one of the enzymes may compensate for loss of the other. To test this, we generated overexpression constructs of *estG* and *bglX,* placed them in a genetic background lacking the other gene, induced overexpression, and subjected the strains to ampicillin treatment. Overproduction of BglX in a Δ*estG* background completely rescued the β-lactam sensitivity of Δ*estG* (Figure 4D). Surprisingly, the reverse was not true—overproduction of EstG did not compensate for loss of *ΔbglX*, which was still sensitive to ampicillin (Figure 4D). Therefore, though there is a genetic interaction between *estG* and *bglX*, these results suggest that EstG and BglX are not functionally redundant.

### *ΔestG* and Δ*bglX* sensitivities are similar to OPG deficient mutants

Inspired by the genetic links to *bglX* and *opgH* that implicated *estG* in the OPG pathway, we wondered whether other aspects of the Δ*estG* phenotype align with the behavior of OPG mutants in other bacteria. In *P. aeruginosa*, OPG production is important for resistance to the ribosome-targeting aminoglycoside antibiotics (Bontemps-Gallo and Lacroix, 2015). Indeed, we found that Δ*estG*, Δ*bglX*, and Δ*estGΔbglX* all have decreased MIC values when treated with the ribosome-targeting antibiotics spectinomycin or tetracycline (Figure 2D). We hypothesized that, like OPG mutants, deletion of *estG* or *bglX* creates a general disruption of the cell envelope, allowing antibiotics to more easily enter the cell, resulting in lower MIC values.

In *E. coli,* OPG synthesis mutants demonstrate increased sensitivity to outer membrane detergents (Rajagopal et al., 2003). We therefore assessed *estG* and *bglX* mutants for sensitivity to the detergent sodium deoxycholate (NaDOC). At 0.6 mg/mL, NaDOC impaired growth of Δ*estG* and Δ*bglX* mutants, and almost entirely inhibited growth of the double mutant (Figure 4—Figure supplement 1C). The sensitivities of Δ*estG* and/or Δ*bglX* strains to ribosome-targeting antibiotics and detergents are consistent with a putative role for both EstG and BglX in maintaining cell envelope integrity via the OPG pathway.

### Δ*estG* sensitivities are rescued by increasing osmolarity

In some organisms, increased OPG production is thought to compensate for a decrease in environmental osmolarity. In low osmolarity media, OPGs in *E. coli* comprise up to 5% of the dry weight, while in high osmolarity media, OPGs account for as low as 0.5% of the dry weight (Bontemps-Gallo et al., 2017). With our hypothesis that Δ*estG* and Δ*bglX* are defective at some point in the OPG pathway, we altered media osmolarity to assess reliance on OPGs in our mutants. We tested this by adding solutes to the media to increase the osmolarity, which we predicted would alleviate the sensitivities of Δ*estG* and Δ*bglX*. When grown in complex media (peptone yeast extract (PYE)), Δ*estG, ΔbglX,* and *ΔestGΔbglX* are all hypersensitive to 50 µg/mL ampicillin (Figure 4E). However, these sensitivities are almost completely alleviated when PYE + ampicillin is supplemented with 50 mM Tris-HCl to increase the osmolarity (Figure 4E). The change in osmolarity does not rescue all mutants with ampicillin sensitivity, as we do not see rescue for a strain bearing deletion of the primary ý-lactamase, *blaA* (West et al., 2002). We see a similar result when sodium chloride is provided as an osmolyte instead of Tris-HCl (Figure 4— Figure supplement 1D). This osmolarity-dependent rescue is further evidence supporting a link between EstG, BglX, and OPGs, and led us hypothesize that EstG acts on OPGs.

### EstG structurally resembles and functions as an esterase *in vitro*

To obtain more insight into a putative substrate for EstG, we determined its structure to 2.1 Å resolution using X-ray crystallography (Figure 5A, PDB ID 7UIC). The EstG final map shows well-defined density for amino acids 30 to 352 and 367 to 444 with excellent geometry (Figure 5A, Table 2). EstG is annotated as a member of the transpeptidase superfamily, and within this family are the well-studied PG enzymes with an α/β hydrolase fold, such as penicillin binding proteins (PBPs) and carboxypeptidases. EstG displays a seven stranded, antiparallel β-sheet sandwiched by the N- and the C-terminal helices in the front and other helices in the back (Figure 5A). The hydrolase domain in EstG is formed by amino acids 30 to 121 and 218 to 444 and displays two motifs that are highly conserved (Ryu et al., 2016). Motif I consists of the Ser-X-X-Lys sequence, residues 101-104, in EstG (Figure 5B) located at the beginning of helix α2, similar to the structure of EstB, a cytoplasmic esterase from *Burkholderia gladioli* (PDB ID 1CI8, Figure 5—Figure supplement 1). Motif II contains a highly conserved Tyr, which acts as a base to activate the serine nucleophile. In EstG, this is Tyr218 (Figure 5B, Figure 5—Figure supplement 1) and is also conserved in the other proteins that share this same fold (Figure 5—Figure supplement 1). Motif I and II are both located in the active site at about 2.7 Å from each other (Figure 5A and B).

**Figure 5:**
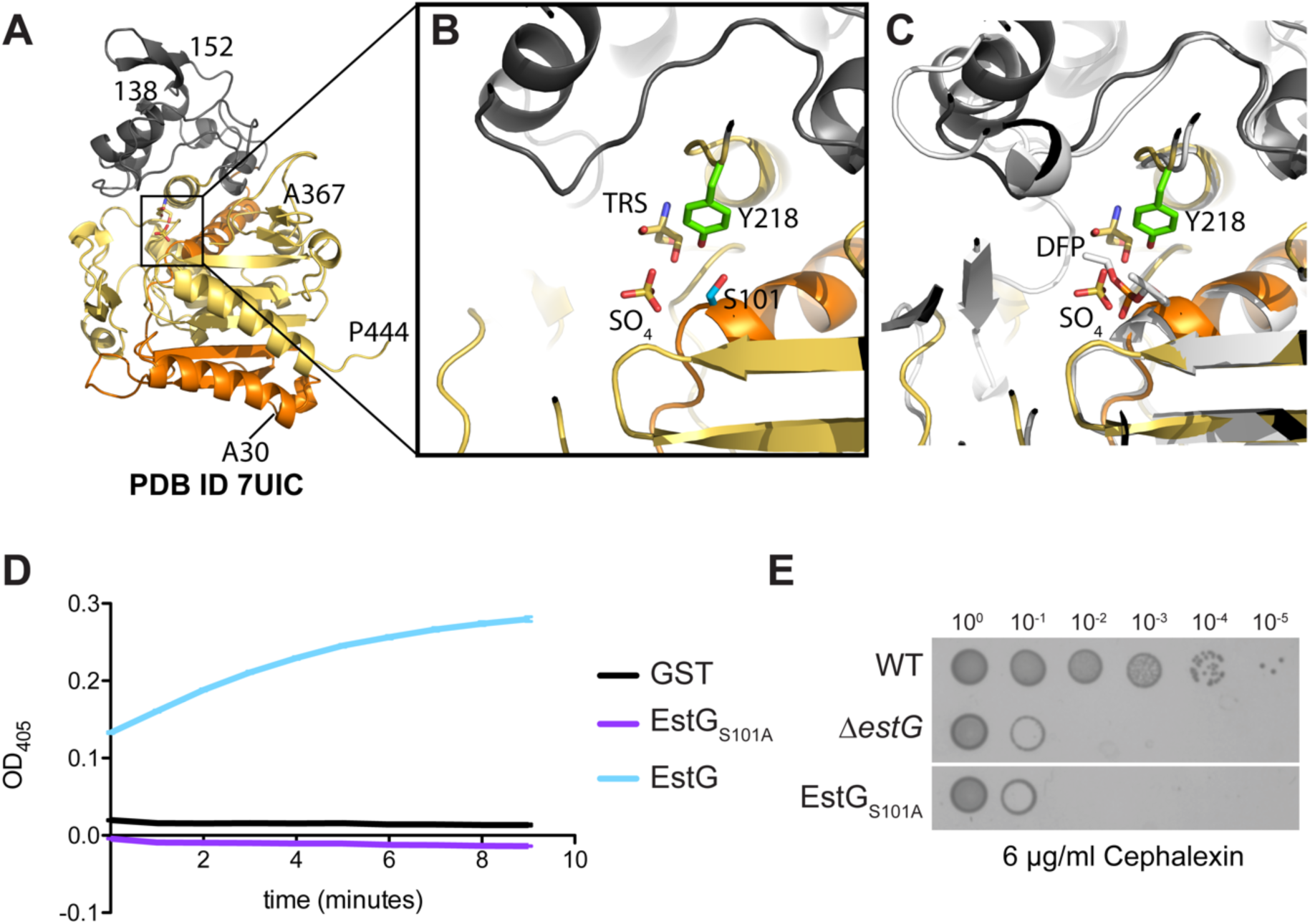
EstG is structurally similar to esterases in the β-lactamase family. **A.** The structure of EstG displays an α/β hydrolase fold. Ribbon diagram of residues 30-444 with the N-terminal residues 30 to 121 colored in orange, 122-217 colored grey, 218 to 444 in yellow. **B.** Zoom in of putative active site identified by homology to esterases. Ser101 (S101) of motif I is 2.7 Å away from Tyr218 (Y218) of motif II. The active site has a sulfate (SO_4_) and a Tris (TRS) molecule bound. **C.** The structural alignment of EstG+ TRS + SO_4_ (PDB ID 7UIC) with EstB bound to diisopropyl fluorophosphate (DFP) (PDB ID 1CI8 (Wagner et al., 2009), colored in light grey) displays the partial overlap of the sulfate to the phosphonate of DFP. **D.** p-nitrophenyl butyrate (pNB) hydrolysis of purified EstG, EstG_S101A_, and GST at 10 µM measured at OD_405_. **E.** Spot dilutions of WT (EG865), Δ*estG* (EG2658), and EstG_S101A_ (EG2990) on PYE agar plates with 6 µg/mL cephalexin. Culture dilutions are as indicated.

**Table 2.** X-ray crystallography data collection and refinement statistics.

|  | **EstG+TRS**<br>**(PDB ID 7UDA)** | **EstG+SO<sub>4</sub>+TRS**<br>**(PDB ID 7UIC)** | **EstG+(Ta<sub>6</sub>Br<sub>12</sub>)<sup>2</sup>**<br>**(PDB ID 7UIB)** |
| --- | --- | --- | --- |
| **Data Collection** | May 27, 1v | Jul 16 (AMX), Au | Jul 16 (AMX) |
| Diffraction source | NSLS-II X17-ID-2 | NSLS-II X17-ID-1 | NSLS-II X17-ID-1 |
| Wavelength (Å) | 0.979321 | 0.920120 | 0.920120 |
| Temperature (K) | 100 | 100 | 100 |
| Detector | Dectris EIGER X 16M | Dectris EIGER X 9M | Dectris EIGER X 9M |
| Space group | P4<sub>1</sub> | P4<sub>1</sub> | P4<sub>1</sub> |
| *a*, *b*, *c* (Å) | 110.3, 110.3, 55.9 | 111.2, 111.2, 56.8 | 111.7, 111.7, 57.4 |
| $\alpha$, $\beta$, $\gamma$ (°) | 90.0, 90.0, 90.0 | 90.0, 90.0, 90.0 | 90.0, 90.0, 90.0 |
| Resolution range (Å) | 29.60–2.47<br>(2.57–2.47) | 28.44–2.09<br>(2.14–2.09) | 28.71–2.62<br>(2.75–2.62) |
| Total no. of reflections | 162,539 (16,345) | 288,425 (19,596) | 195,511 (22,742) |
| No. of unique reflections | 24,085 (2,515) | 41,361(2,900) | 21,379(2,741) |
| Completeness (%) | 99.0 (91.9) | 99.0 (91.9) | 99.5 (96.9) |
| Redundancy | 6.7 (6.5) | 7.0 (6.8) | 9.1(8.3) |
| $\langle I/\sigma(I) \rangle$ | 10.9 (2.4) | 12.4 (2.6) | 16.9(3.1) |
| R<sub>merge</sub> | 0.99 (0.71) | 0.086 (0.77) | 0.10(0.76) |
| R<sub>meas</sub> | 0.11 (0.84) | 0.10 (0.90) | 0.11(0.81) |
| R<sub>pim</sub> | 0.06 (0.44) | 0.05(0.47) | 0.03(0.27) |
| CC<sub>1/2</sub> | 0.99 (0.75) | 0.99 (86.3) | 0.99(0.79) |
| **Refinement** |  |  |  |
| Resolution range (Å) | 29.62–2.47<br>(2.53–2.47) | 27.83–2.09<br>(2.14–2.09) | 27.96–2.70<br>(2.77–2.70) |
| No. of reflections, working set | 22,895,1171 | 39,246 | 18,687 |
| R<sub>work</sub>/R<sub>free</sub> | 0.18/0.22<br>(0.30/0.35) | 0.17/0.21<br>(0.24/0.25) | 0.20/0.24<br>(0.30/0.32) |
| *No. of non-H atoms* |  |  |  |
| EstG | 3,093 | 3,031 | 3069 |
| ligand | 34 | 7 | 39 |
| Total of non-H atoms | 3,127 | 3,253 | 3,085 |
| *R.m.s. deviations* |  |  |  |
| Bonds (Å) | 0.009 | 0.012 | 0.012 |
| Angles (°) | 1.74 | 0.001 | 1.731 |
| Wilson B-factor (Å<sup>2</sup>) | 52 | 43 | 47 |
| *Average B factors (Å<sup>2</sup>)* |  |  |  |
| EstG | 54 | 42 | 52 |
| ligand | 58 | 43 | 80 |
| Total average B factor | 56 | 42 | 68 |

| *Ramachandran (%)* |  |  |  |
| --- | --- | --- | --- |
| Favorable | 95.2 | 96.98 | 94.36 |
| Allowed | 3.8 | 2.26 | 3.92 |
| Outlier | 1.0 | 0.76 | 1.72 |

\*Values in parentheses are for highest-resolution shell. All atoms refer to non-H atoms.

In total, we determined three structures of EstG: EstG bound to tris (EstG+TRS), EstG bound to tris and sulfate (EstG+TRS+SO_4_), and EstG bound to tris, sulfate, and tantalum bromide (EstG+TRS+SO_4_+(Ta_6_Br_12_)^2^). The structures are very similar with a pairwise root-mean-square deviation ranging 0.24 to 0.26 Å for amino acids 398-401 as calculated with SSM Coot (Emsley and Cowtan, 2004). The binding of a SO_4_ molecule close to Motif I and Motif II correlates with the presence of clear electron density for the loop 346-357 (PDB ID 7UIC, 7UIB, Figure 5B, Figure 5—Figure Supplement 2). Structural alignment of EstG+TRS+SO_4_ with EstB bound to diisopropyl fluorophosphate (DFP, PDB ID 1CI9) highlights the partial overlap between the SO_4_ in EstG and the DFP bound to catalytic serine residue in EstB (Figure 5C).

In EstG, residues 122 to 217 are on top of the hydrolase fold (Figure 5A). Within it, residues 138 through 151 define an insertion of a hairpin formed by strand β4-β5 (Figure 5--Figure supplement 1, 3) which is also present in the transesterase enzyme, simvastatin synthase (*Aspergillus terreus* LovD, PDB ID 4LCM). Notably, this hairpin is absent in EstB.

The structural alignment over those deposited in the PDB highlights a structural conservation among enzymes in this large family of proteins. The most similar structures to EstG by structure/sequence are esterases (EstB, PDB ID 1CI8), transesterases (LovD, PDB ID 3HLB), carboxylesterases (PDB ID 4IVK), PBP homologs (PDB ID 2QMI), and D-amino acid amidases (PDB ID 2DNS). All these enzyme classes are referenced in the literature as having homology to β-lactamase folding esterases (Ryu et al., 2016). Interestingly, D-amino acid amidases and aminohydrolases also lack the hairpin insertion described for EstG (Figure 5—Figure Supplement 3).

Based on the structural similarity of EstG to EstB, a cytoplasmic esterase with an unknown native substrate, we sought to compare the two enzymatically. Despite the β-lactamase fold, EstB has no β-lactamase or peptidase activity (Wagner et al., 2009), similar to our observations with EstG (Figure 2—Figure supplement 1E, 2). EstB does, however, demonstrate esterase hydrolytic activity (Wagner et al., 2009). *In vitro* esterase activity can be detected using p-nitrophenyl esters, such as p-nitrophenyl butyrate (pNB), as hydrolysis of the substrate creates a visible color change that can be measured as absorbance over time (similar to pNPG hydrolysis). Using this assay, EstG significantly hydrolyzed pNB as compared to the negative control, GST (Figure 5D). We sought to create a catalytically dead mutant of EstG by mutating the predicted active site serine, Ser101, within motif I. Consistent with our prediction, the S101A mutant cannot hydrolyze pNB *in vitro*, confirming that it is a catalytically dead variant (Figure 5D). Additionally, when Ser101 is mutated to alanine (S101A) in the chromosomal copy of *estG*, this mutant phenocopies the β-lactam sensitivity of Δ*estG in vivo* (Figure 5E). These data establish the essentiality of EstG’s enzymatic activity in protecting the cell against stress and confirms activity of EstG as an esterase.

### EstG enzymatically modifies a cyclic hexasaccharide periplasmic glucan

EstG can act as an esterase *in vitro* and our genetic and osmolarity data implicate OPGs as a substrate. However, *Caulobacter* OPGs have never been characterized, and the absence of homologs of most characterized OPG-metabolizing enzymes in this organism precludes a simple prediction of which OPG species may be present. To identify the native substrate of EstG, we fractionated WT cells into periplast and spheroplast fractions followed by isolation of putative periplasmic sugars (Figure 6A). Given that *E. coli* OPGs are between 1 to 10 kDa, we hypothesized that *Caulobacter* OPGs might be of similar size. Therefore, we further fractionated to isolate only components within our desired size range. The remaining sample was boiled to remove contaminating proteins, leaving sugars or other heat-resistant metabolites intact. *In vitro,* we combined this 1-10 kDa periplast isolate with purified EstG or the catalytically dead mutant, EstG_S101A_. We then separated molecules in the treated periplast by high-performance liquid chromatography (HPLC) and selected for peaks that decreased in abundance when mixed with EstG, but not when mixed with EstG_S101A_. Peaks of interest were then identified by mass spectrometry. Using this approach, we identified a molecule that decreased in abundance ∼40% when incubated with EstG (Figure 6B), indicating that EstG enzymatically modified this substrate in some way. The mass of the parental ion led us to hypothesize that the molecule resembled α-cyclodextrin (α-CD), a cyclic, hexameric glucose polymer. Notably, the MS/MS spectra for this molecule in the periplast + EstG_S101A_ (top half of Figure 6C), most closely matches the library spectra for α-CD (bottom half of Figure 6C). Greater than 80% of the fragmentation signal generated from our experiments match the ion profile for α-CD. We next attempted to detect chemical modification of α-CD by EstG using our periplast and mass spectrometry workflow. However, due to the complexity of the periplast fraction and the small expected amount of modified α-CD, we were not able to identify a modified α-CD molecule or determine a specific activity of EstG on α-CD. Though this small, cyclic sugar is a novel structure for an OPG, it is consistent with the existence of cyclic OPGs in other bacteria, notably Family IV cyclic OPGs synthesized by OpgH in *Rhodobacter sphaeroides* and related α-proteobacteria (Bontemps-Gallo et al., 2017).

**Figure 6:**
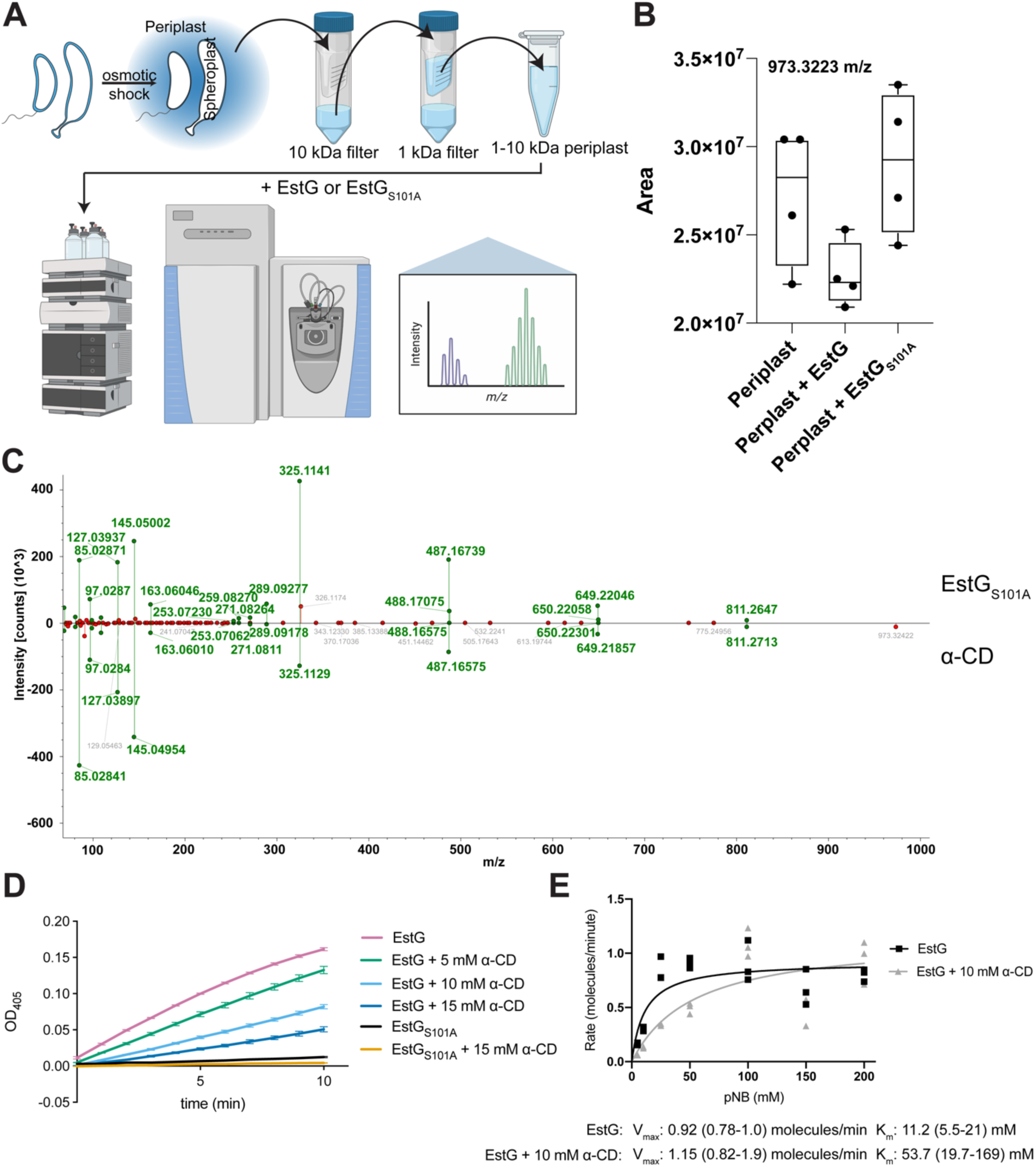
A cyclic hexameric glucose is the native substrate of EstG. **A.** Schematic outlining the method for isolating periplasmic contents (periplast, blue) and sequential fractionation. Periplast 1-10 kDa was then combined with EstG or EstG_S101A_, and contents were separated and identified with LCMS. **B.** A box-plot displaying the relative abundances of the cyclic hexaglycan with error bars across four technical replicates in the samples of periplast alone, periplast + EstG, or periplast + EstG_S101A_. Mass of the parental ion is 973.3223 m/z. **C.** MS/MS spectra with the experimental spectra observed in one of the injections of the periplast + EstG_S101A_ as the top of the mirror plot. Bottom half of the mirror plot is the mzCloud reference spectra for α-cyclodextrin (α-CD). **D.** p-nitrophenyl butyrate (pNB) hydrolysis of purified EstG or EstG_S101A_ with increasing amounts of α-CD showing concentration dependent inhibition. **E.** Michaelis–Menten saturation curve of the rates of pNB hydrolysis with EstG or EstG + 10 mM α-CD to show competitive inhibition of the active site. Rate was determined by the slope of the pNB hydrolysis curve at the indicated pNB concentration. Rate is presented as molecules of pNB hydrolyzed per minute. Parenthesis next to values for V_max_ and K_m_ represent 95% confidence interval.

We next sought to validate α-CD as an EstG substrate *in vitro*. If α-CD is a substrate for EstG, we reasoned we could add α-CD to the pNB hydrolysis assay and inhibit pNB hydrolysis through competition for the active site. Indeed, increasing amounts of α-CD reduced EstG’s hydrolysis of pNB in a concentration-dependent manner, while EstG_S101A_ remained unchanged with added α-CD (Figure 6D). Though the inhibition is clearly concentration-dependent, we wanted to confirm that α-CD was competitively inhibiting EstG’s active site, consistent with it being a substrate. To achieve this, we measured the rate of pNB hydrolysis with increasing concentrations of pNB and a consistent amount of α-CD. For a competitive inhibitor, we expect to see a constant V_max_ and an increased K_m_ value with added α-CD. By plotting the rate of hydrolysis +/- α-CD, the V_max_ values of the two curves are close at 0.92 molecules/min without α-CD and 1.15 molecules/minute with α-CD (Figure 6E, Figure 6—Figure supplement 1). However, the K_m_ values differ, at 11.2 mM without α-CD and 53.7 mM with α-CD (Figure 6E, Figure 6—Figure supplement 1). These values produce the expected pattern for a competitive inhibitor and gave us confidence that α-CD interacts directly with the active site of EstG and is thus structurally similar to the native substrate. Collectively, these data suggest that EstG modifies a previously uncharacterized cyclic, hexameric OPG in a novel manner, thereby contributing to cell envelope homeostasis during stress (Figure 7).

**Figure 7:**
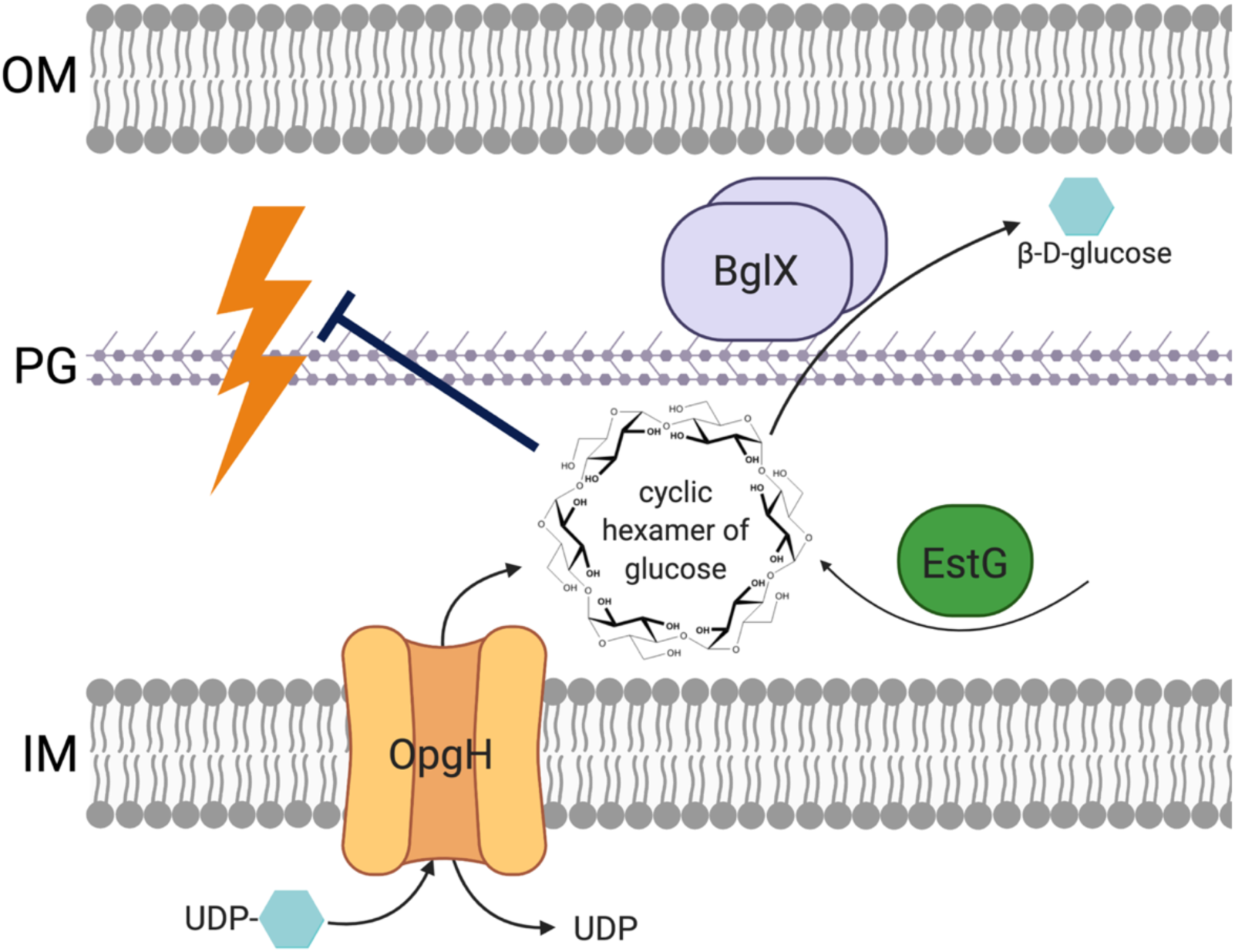
EstG protects the cell envelope against stress through its activity on cyclic OPG polymers. Cell envelope homeostasis during stress is maintained through the actions of EstG and the putative OPG pathway in *Caulobacter*. We propose that OpgH takes cytoplasmic UDP-glucose to synthesize small, cyclic OPG molecules into the periplasm. We believe BglX hydrolyzes these OPGs and EstG modifies it in some way to modulate the osmolarity of the periplasm. Without OPG production or modification, the cell envelope integrity is lost, resulting in hypersensitivity to a variety of environmental changes and antibiotic stresses (represented by yellow lightning bolt). OM=outer membrane, PG=peptidoglycan, IM=inner membrane.

## Discussion

It is clear from our work that proteins and pathways that play critical roles in maintaining cell envelope homeostasis remain undiscovered. Our identification of EstG and its novel role in the *Caulobacter* OPG pathway suggests there might be unexplored substrates of other TPase family enzymes. We identified EstG through a Tn-Seq screen as an essential factor for surviving ΔCTL-induced cell wall stress (Figure 1). Though *estG* is non-essential in unstressed conditions (Figure 2), Δ*estG* is hypersensitive to cell envelope stresses (Figure 2D, Figure 4—Figure supplement 1C). Despite its homology to TPase family proteins, EstG does not detectably modify the PG (Figure 2—figure supplement 2, Table 1). Instead, genetic interactions with *opgH* (Figure 3), encoding the predicted OPG synthase, and *bglX* (Figure 4), encoding a putative OPG hydrolase, implicate EstG in the OPG pathway. *In vitro* biochemistry revealed a periplasmic substrate of EstG as a cyclic hexamer of glucose, which is the first reported OPG in *Caulobacter* (Figure 6).

In this study, we originally set out to identify mechanisms of ΔCTL suppression. We were surprised to primarily recover suppressing mutations in stress response pathways, instead of cell envelope- or cell wall-related genes. Activation of stress response pathways typically leads to sweeping changes in cellular physiology, suggesting that the stress imposed by ΔCTL is multifaceted and cannot easily be suppressed by mutation of a single factor. We leveraged (p)ppGpp-mediated suppression of ΔCTL to identify more direct factors involved in surviving ΔCTL-induced stress and, through this approach, found *estG*. While following up on the role of EstG in (p)ppGpp-dependent suppression of ΔCTL, we found that *estG* is unrelated to (p)ppGpp. Instead, it was the additional antibiotic stress (e.g., introduction of gentamycin marked *relA’* to produce high (p)ppGpp) in the presence of ΔCTL stress that made *estG* essential (data not shown). We further confirmed this by deleting *estG* in a ΔCTL background suppressed by high (p)ppGpp through a hyperactive *spoT* mutant, which was not lethal (data not shown). In retrospect, this finding is not entirely surprising given the critical role we established for EstG in surviving a variety of antibiotic stresses.

Both our own characterization of the Δ*estG* strain and information in the Fitness Browser database (Price et al., 2018; Wetmore et al., 2015) indicated a wide range of antibiotic sensitivities. Those we tested (Figure 2D) include many classes of PG- and ribosome-targeting antibiotics such as β-lactam (ampicillin, mecillinam, cefaphlexin), glycopeptide (vancomycin), phosphonic (fosfomycin), aminoglycoside (spectinomycin), and tetracycline antibiotics (Figure 2D). The Fitness Browser additionally indicated sensitivities to a DNA-gyrase-targeting antibiotic (nalidixic acid) and an inhibitor of lipid A biosynthesis (CHIR-090). Collectively, this establishes Δ*estG* hypersensitivity to antibiotics that target at least four different cellular processes (PG metabolism, protein synthesis, DNA topology, and outer membrane biosynthesis/homeostasis). We looked for similarities among these drug classes but found no obvious biochemical similarities. For instance, nalidixic acid and ampicillin are relatively small, while vancomycin is a large glycopeptide, and though most molecules tested were polar and uncharged, others, such as chloramphenicol (data not shown) and sodium deoxycholate are charged. Ultimately, these broad antibiotic sensitivities support the idea of a global cell envelope defect resulting from loss of EstG’s enzymatic activity, and not a sensitivity specific to a particular molecular feature.

EstG is classified as a ý-lactamase family protein within the TPase superfamily, which is why it stood out as an attractive candidate from a cell wall stress screen. Of characterized proteins, EstG shares the most structural and biochemical similarities to EstB from *B. gladioli*, another enzyme in the ý-lactamase family that adopts an α/ý hydrolase fold. They both contain an active site serine, but EstG lacks the common esterase motif, G-X-S-X-G, present in EstB. This esterase motif is not required for EstB’s hydrolase activity, however (Wagner et al., 2009). Our data provide evidence of EstG acting as an esterase and not a ý-lactamase, and we have also identified a novel EstG substrate. Within *Caulobacter*, EstG is one of eight enzymes that are classified as putative ý-lactamases (West et al., 2002) that potentially do not function as ý-lactamases at all. EstG is just one example of the numerous enzymes across bacteria that fall into the TPase superfamily but have novel activities or substrates.

Though not required under normal growth, our data demonstrate the importance of EstG acting on its sugar substrate and implicates an essential role for OPGs in stress survival. OPGs have not been previously identified in *Caulobacter*, though the presence of an *opgH/mdoH* homolog in the genome was reported (Bohin, 2000). OPGs in several α-proteobacterial species of the orders Rhizobiales and Rhodobacterales are characterized and have a wide variety of sizes and structures, consisting of family II, III, and IV OPGs (Bohin, 2000). These OPGs can range from 10-25 glucose monomers, but all three classes are cyclic polymers, as opposed to the linear family I OPGs commonly found in γ-proteobacteria. We were surprised to find that the only *opg* gene in *Caulobacter* is *opgH*. As we report a cyclic OPG-like molecule, we would expect other OPG genes responsible for cyclizing and modifying OPGs to be present in *Caulobacter*. Uniquely, other α-proteobacteria encode OPG metabolic enzymes that are not homologs of the *opg* genes in *E. coli* including *chvA* and *chvB* in *Agrobacterium tumefaciens*, *ndvA* and *ndvB* in *Sinorhizobium meliloti,* and *cgs* and *cgt* in *Brucella abortus* (Bontemps-Gallo et al., 2017). Distinct from the well-described *opg*/*mdo* genes, these genes imply the existence of a wide variety of OPG enzymes and OPG structures across bacteria. Additionally, among these OPG metabolic genes, there are proteins whose precise enzymatic functions remain elusive, such as NdvD in *S. meliloti* (Bontemps-Gallo et al., 2017). We propose that EstG and BglX are additional examples of enzymes with unique roles in OPG synthesis, modification, and/or hydrolysis.

Mutants of OPG enzymes in diverse bacteria typically have pleiotropic phenotypes, including those discussed for *estG* and *bglX* mutants (e.g. antibiotic sensitivity) as well as defects in motility, biofilm formation, and/or virulence (Bontemps-Gallo et al., 2017). Despite the impact of OPGs on important cellular behaviors and properties, we do not know the mechanism(s) behind OPG- mediated effects. One model suggests OPGs function as osmoprotectants by establishing a Donnan equilibrium across the outer membrane. The idea is that production of negatively charged OPGs in the periplasm (as occurs in *E. coli*) creates a high concentration of fixed, charged molecules that cannot cross the outer membrane. The accumulation of charged OPGs attracts counterions to the periplasm, and maintains a Donnan membrane potential across the outer membrane, allowing for isosmolarity of the periplasm and cytoplasm even in low osmolarity environments (Kennedy et al., 1982; Stock et al., 1977). The Donnan potential has also been suggested to play a role in permeability of the envelope to antibiotics (Alegun et al., 2021). These mechanisms, however, presume that OPGs are always highly charged, which is not the case in all bacteria, and may or may not be the case in *Caulobacter* (Bontemps-Gallo et al., 2017). Though we were not able to determine the exact EstG-mediated modification on *Caulobacter* OPGs, it is possible that EstG adds a charged moiety in order to mediate the Donnan potential and protect the cell envelope.

Beyond the Donnan potential, OPGs are postulated to have other functions in cell envelope homeostasis, such as a role in envelope organization, cell signaling, and protein folding (Bontemps-Gallo et al., 2017). For instance, loss of OPGs in *E. coli* was reported to cause an increase in the periplasmic space of plasmolyzed cells, perhaps reflecting a structural role in maintaining envelope geometry (Holtje et al., 1988). Deletion of *estG*, however, did not result in a notable increase in periplasmic space (data not shown) and suggests that *Caulobacter opg* mutants may not directly impact the structure of the periplasm.

Despite the unclear mechanism of OPG-mediated envelope protection, our data suggest that the modification and/or hydrolysis activity of EstG and BglX on *Caulobacter* OPGs contributes to osmoprotective properties, most notably supported by the osmolarity-dependent rescue of antibiotic sensitivity (Figure 4E). It is possible that more mechanistic insight can be revealed with further study of OPG pathways and enzymes in other organisms. For instance, the *E. coli* OpgH enzyme links nutrient availability with cell size by inhibiting FtsZ when UDP-glucose levels are high (Hill et al., 2013). This is likely not a conserved function of *Caulobacter* OpgH, as it lacks most of the N-terminal FtsZ-interacting region. Suppressor mutations within *opgH* have also been identified in *E. coli* that further implicate OpgH with envelope homeostasis. A nonsense mutation in *opgH* was isolated in a lipopolysaccharide (LPS) mutant that together conferred resistance to a polypeptide antibiotic (bacitracin), a polyketide antibiotic (rifampin), and sodium dodecyl sulfate (Falchi et al., 2018). Due to the integral role of LPS in outer membrane integrity, it was proposed that either the lack of OPGs or loss of OpgH reduces membrane permeability to antibiotics, thus conferring resistance. However, unlike the *opgH* suppressing mutations identified in this study (L480P and L434), the *E. coli opgH* nonsense mutation was recessive to WT. Though this suggests a different mechanism of suppression, it does not rule out the possibility of deficient OPG production in the *Caulobacter opgH* mutants, resulting in a less permeable membrane and our observed resistance to stress. Two spontaneous *opgH* mutants were also isolated in *Vibrio cholerae* that suppressed the hyperosmotic lethality of a lytic transglycosylase (LTG) mutant (Weaver et al., 2022). This model suggested that LTG mutants inadequately recycle PG products, resulting in excessive periplasmic crowding (Weaver et al., 2022). Additional production of OPGs exacerbated this periplasmic crowding, which was lethal in low osmolarity environments (Weaver et al., 2022). Though the hyperosmotic growth defect of LTG mutants and periplasmic crowding could indicate a similar role for EstG, the identification of an OPG substrate indicates a direct link to OPG metabolism, rather than an indirect consequence of molecular crowding. An important avenue for future work includes functional studies of OpgH and these mutants as well as determination of the exact structure and potential modifications on *Caulobacter* OPGs. These insights can ultimately bridge our gap in understanding of the mechanistic role of OPGs in the *Caulobacter* envelope.

## Materials and methods

### Caulobacter crescentus and Escherichia coli growth media and conditions

*C. crescentus* NA1000 cells were grown at 30°C in peptone-yeast extract (PYE) medium. *E. coli* Rosetta(DE3)/pLysS cells were grown at 30°C in Luria-Bertani (LB) medium. Xylose or glucose were used at concentrations of 0.3% (w/v) for induction experiments. Antibiotics were used in liquid (solid) medium at the following concentrations for *Caulobacter* growth: gentamycin, 1 (5) µg/mL; kanamycin, 5 (25) µg/mL; spectinomycin, 25 (100) µg/mL. Streptomycin was used at 5 µg/mL in solid medium. *E. coli* antibiotics were used in liquid (solid) medium as follows: ampicillin, 50 (100) µg/mL; gentamicin, 15 (20) µg/mL; kanamycin, 30 (50) µg/mL; and spectinomycin, 50 (50) µg/mL. For growth curves, a Tecan Infinite M200 Pro plate reader measured absorbance every 30 minutes at OD_600_ of a 100 µL culture volume in a 96 well plate in biological triplicate with intermittent shaking. For spot dilution assays, mid-log cells were diluted to an OD_600_ of 0.05 and serially diluted up to 10^-6^ before spotting 5 µL of each dilution onto a PYE plate with indicated inducer and/or antibiotic. Plates were incubated at 30°C for 48 hours, or until the appearance of colonies at the lowest dilution in the control strain. To determine the minimum inhibitory concentration (MIC), mid-log phase cells were diluted to OD_600_ of 0.5 and 200 µL were spread out onto a PYE plate. Antibiotic strips with increasing concentration of antibiotic were placed on the dried plate, inverted, and grown at 30°C for 48 hours. Some MIC values were estimated by loss of growth on plates with a range of antibiotic added to the media. A summary of all strains, plasmids, and primers used in this study can be found in Supplement Table 3.

### Atypical strain construction

We were unable to generate the following strains in low osmolarity PYE media, so they were constructed in M2G minimal media: EG3116 (ΔCTL+Δ*estG*), EG3369 (*opgH_L480P_*), EG3371 *(ΔestG*+OpgH_L480P_), and EG3377 (*P_van_-opgH*). For a 500 mL batch of M2G plates, 465 mL of water and 7.5 g agar (1.5%) were autoclaved. Once cooled, 25 mL of 5x M2 salts, 500 µL of 500 mM MgSO_4_, 500 µL of 10 mM FeSO_4_ 10 mM EDTA (Sigma F-0518), and 0.3% glucose were added. Additional antibiotics or media supplements needed for selection were also added at this time.

### Phase-contrast microscopy

Exponential phase cells were spotted on 1% agarose pads and imaged using a Nikon Eclipse Ti inverted microscope equipped with a Nikon Plan Fluor 100X (NA1.30) oil Ph3 objective and Photometrics CoolSNAP HQ^2^ cooled CCD camera. Images were processed using Adobe Photoshop.

### Suppressor screening and whole genome sequencing

For the ΔCTL suppressor screen, *Caulobacter* strains EG937 or EG1214 strains were inoculated from individual colonies and grown overnight in PYE media (with no inducer) until stationary phase. Cells were plated on PYE agar plates containing 0.3% (w/v) xylose to induce ΔCTL expression and incubated at 30°C until the appearance of colonies (suppressors). Suppressors were tested for growth in PYE media with 0.3% xylose overnight. Immunoblotting with FtsZ- antiserum was used to confirm xylose-induced ΔCTL expression. Genomic DNA was extracted from suppressors using Qiagen DNeasy Blood and Tissue Kit. Mutations were identified from MiSeq analysis of genomic DNA from suppressor strains. Spontaneous suppressors of Δ*estG* were isolated by plating Δ*estG* (EG2658) on PYE+100 µg/mL ampicillin and isolating resistant colonies. Resistance was confirmed by spot dilution on plates containing 50 µg/mL ampicillin. Genomic DNA was extracted from suppressors using Qiagen DNeasy Blood and Tissue Kit and sent to Microbial Genome Sequencing Center (MiGS) for whole genome sequencing and BreSeq analysis.

### Cell fractionation

Cells were fractionated into periplasm and spheroplast using the previously described methods in Judd et al, except that 2 µg/mL lysozyme was used (Judd et al., 2005). Briefly, cells were grown at 30° to an OD_600_ of 0.5 in 10 mL of PYE. Cells were pelleted at 3,500 x g for 10 minutes and the supernatant removed. The pellet was resuspended in 1 mL of periplasting buffer (50 mM Tris-HCl pH 8.0, 18% sucrose, and 1 mM CaCl_2_) and then 2 µg/mL of lysozyme and 1 mM EDTA was added. Contents were left on ice for 30 minutes and then spun at 3,140 x g for 5 minutes. The supernatant (periplast fraction) was carefully removed to a fresh tube, and the pellet (spheroplast fraction) was saved.

### Transposon library preparation, sequencing, and analysis

Transposon libraries were prepared, sequenced, and analyzed using the same methods as previously described in Woldemeskel et al. and Lariviere et al. (Woldemeskel et al., 2020; Lariviere et al., 2019). Tn-Seq libraries were generated for WT (EG865), RelA’ (EG1799) and ΔCTL+RelA’ (EG1616). 1L PYE cultures were harvested at OD_600_ of 0.4–0.6, washed 5 times with 10% glycerol, and electroporated with the Ez-Tn5 <KAN-2> transposome (Epicentre, Charlotte, North Carolina). Cells recovered at 30°C shaking for 90 minutes, and plated on PYE-Kan plates. The RelA’ library was plated on PYE-Kan with gentamycin and 0.003% xylose to induce RelA’ expression. ΔCTL+RelA’ library was plated on PYE-Kan plates with spectinomycin, streptomycin, gentamycin, and 0.003% xylose to induce RelA’ and ΔCTL. Colonies were scraped off plates, combined, resuspended to form a homogeneous solution in PYE, and flash frozen in 20% glycerol. The DNeasy Blood and Tissue Kit (Qiagen, Hilden, Germany) was used to extract genomic DNA from each pooled library. Libraries were prepared for Illumina Next-Generation sequencing through sequential PCR reactions. The initial PCR round used arbitrary hexamer primers with a Tn5 specific primer going outward. The second round used indexing primers with unique identifiers to filter artifacts arising from PCR duplicates. Indexed libraries were pooled and sequenced at the University of Massachusetts Amherst Genomics Core Facility on the NextSeq 550 (Illumina, San Diego, California).

Sequencing reads were first demultiplexed by index, each library was concatenated and clipped of the molecular modifier added in the second PCR using Je (Girardot et al., 2016):

java -jar /je_1.2/je_1.2_bundle.jar clip F1 = compiled.gz LEN = 6

Reads were then mapped back to the Caulobacter crescentus NA1000 genome (NCBI Reference Sequence: NC_011916.1) using BWA (Li and Durbin, 2010) and sorted using Samtools (Li et al., 2009):

bwa mem -t2 clipped.gz | samtools sort -@2 - > sorted.bam

Duplicates were removed using Je (Girardot et al., 2016) and indexed with Samtools (Li et al., 2009) using the following command:

java -jar /je_1.2/je_1.2_bundle.jar markdupes I = sorted.bam O = marked.bam M = METRICS.txt MM = 0 REMOVE_DUPLICATES = TRUE

samtools index marked.bam

The 5’ insertion site of each transposon were converted into .wig files comprising counts per position and visualized using Integrative Genomics Viewer (IGV) (Robinson et al., 2011; Thorvaldsdottir et al., 2012). Specific hits for each library were determined with coverage and insertion frequency using a bedfile containing all open reading frames from NC_011916.1 and the outer 20% of each removed to yield a clean and thorough insertion profile. This was determined using BEDTools (Mccarthy et al., 2012; Robinson et al., 2010) and the following commands:

bedtools genomecov -5 -bg marked.bam > marked.bed

bedtools map -a NA1000.txt -b marked.bed -c 4 > output.txt

Tn-Seq data have been deposited in the Sequence Read Archive (SRA) under accession numbers:

### Protein purification

All purified proteins were overproduced in Rosetta (DE3) pLysS *E. coli* from the following plasmids: His_6_-EstG-His_6_, pEG1622; His_6_-EstG_S101A_-His_6_, pEG1706; His_6_-EstA, pEG1950; His_6_-BglX-His_6_, pEG1779. Cells were induced with 1mM IPTG for 4 hours at 30°C. Cell pellets were resuspended in Column Buffer A (50 mM Tris-HCl pH 8.0, 300 mM NaCl, 10% glycerol, 20 mM imidazole, 1 mM β-mercaptoethanol) flash frozen in liquid nitrogen and stored at -80°C. To purify the His-tagged proteins, pellets were thawed at 37°C, and 10 U/mL DNAse 1, 1 µg/mL lysozyme, and 2.5 mM MgCl_2_ were added. Cell slurries were left on ice and occasionally inverted for 45 minutes, then sonicated and centrifuged for 30 minutes at 15,000 x *g* at 4°C. The protein supernatant was then filtered and loaded onto a pre-equilibrated HisTrap FF 1mL column (Cytiva, Marlborough, Massachusetts). The His-tagged proteins were eluted in 30% Column Buffer B (same as Column Buffer A but with 1M imidazole). Peak fractions were concentrated and applied to a Superdex 200 10/300 GL (Cytiva) column equilibrated with EstG storage buffer (50 mM HEPES-NaOH pH 7.2, 150 mM NaCl, 10% glycerol, 1 mM β-mercaptoethanol). Peak fractions were combined, concentrated, and snap-frozen in liquid nitrogen and stored at -80°C.

### Immunoblotting

Purified His_6_-EstG-His_6_ was dialyzed into PBS and used to immunize a rabbit for antibody production (Pocono Rabbit Farm & Laboratory, Canadensis, Pennsylvania). To affinity purify the EstG antisera, His_6_-EstG-His_6_ in EstG storage buffer was coupled to Affigel 10 resin (Bio-Rad, Hercules, California). After washing the resin 3 times with cold water, add approximately 10 mg of protein to 1 mL of Affigel 10 resin to rotate at 4°C for 4 hours. 75 mM Tris pH 8.0 was added to terminate the reaction and left to rotate at 4°C for 30 minutes. EstG-resin was washed in a column with the following cold reagents: 10 mL EstG storage buffer, 15 mL Tris-buffered saline (TBS), 15 mL 0.2 M glycine-HCl pH 2.5 with 150 mM NaCl, 15 mL TBS, 15 mL guanidine-HCl in TBS, and 20 mL TBS. EstG antisera was combined with EstG-resin, and incubated, rotating, overnight at 4°C. Unbound sera flowed through the column and was washed with 25 mL TBS, 25 mL TBS with 500 mM NaCl and 0.2% Triton X-100, and a final wash of 25 mL TBS. Bound Anti-EstG was eluted with 0.2 M glycine pH 2.5 and 150 mM NaCl, dialyzed into TBS, and diluted 1:1 with glycerol. Anti-EstG antibody specificity was validated by western blot to recognize a band in wild type lysate that is absent in a Δ*estG* mutant.

Western blotting was performed using standard lab procedures. Cells in log phase were isolated and lysed in SDS-PAGE loading buffer and boiled for 10 minutes. For a given experiment, equivalent OD units of cell lysate were loaded. SDS-PAGE and transfer of protein to nitrocellulose membrane were performed using standard procedures. Antibodies were used at the following concentrations: EstG-1:1000; SpmX-1:10,000 (Radhakrishnan et al., 2008); Flag-1:1,000 (Sigma, St. Louis, Missouri); CdnL-1:2,500 (Woldemeskel et al., 2020).

### *In vitro* pNB hydrolysis or pNPG assay

To test for serine hydrolase activity using p-nitrophenyl butyrate (pNB, Sigma), indicated proteins were used at 10 µM in a 50 µL reaction containing 50 mM Tris-HCl pH 8. pNB was added last to the samples at a concentration of 4 µM. Absorbance was measured every minute at 405 nm for 10 minutes. To test for glucosidase activity using 4-Nitrophenyl-β-D-glucopyranoside (pNPG, Sigma), indicated proteins were used at listed concentrations in a 50 µL reaction containing 50 mM Tris-HCl pH 8. pNPG was added last at a final concentration of 4 µM. Absorbance was measured every minute at 405 nm for 10 minutes.

### Nitrocefin hydrolysis assay

To assess β-lactamase activity through hydrolysis of nitrocefin, 10 µM of indicated proteins were mixed with 100 µM nitrocefin (Calbiochem, Sigma) in a reaction buffer containing EstG storage buffer (50 mM HEPES-NaOH pH 7.2, 150 mM NaCl, 10% glycerol, 1 mM BME) to a final volume of 100 µL. Absorbance was measured at 492 nm every 10 minutes for 4 hours.

### Sacculi purification and PG binding assay

Sacculi for PG binding assay were prepared as previously described in Meier et al (Meier et al., 2017). Wild type (EG865) *Caulobacter* cells were grown in 1L of PYE at 30°C to an OD_600_ of 0.5. Cells were pelleted by centrifugation at 6,000 x *g* for 10 minutes and resuspended in 10 mL of 1X PBS. The cells were added dropwise to a boiling solution of 4% SDS where they were continuously mixed and boiled for 30 minutes, then incubated overnight at room temperature. Sacculi were pelleted by ultracentrifugation at 42,000 x g in an MLA-80 rotor for 1 hour at 25°C and remaining pellet was washed four times with ultra-pure water with a final resuspension in 1 mL PBS with 20 µL of 10 mg/mL amylase, left at room temperature overnight. Then the sacculi were pelleted at 90,000 x g in an MLA-130 rotor for 15 minutes at 25°C and washed three times with ultra-pure water, with a final resuspension in 1 mL of PG binding buffer (20 mM Tris-HCl pH 6.8, 1 mM MgCl_2_, 30 mM NaCl, 0.05% Triton X-100). To each reaction, 6 µg of each protein was added to either PG or buffer. Reactions were left on ice for 30 minutes and then centrifuged for 30 minutes at 90,000 x g in the MLA-130 rotor at 4°C. Supernatant was saved and the pellet was resuspended in PG binding buffer and saved as the PG bound isolate. SDS-PAGE loading dye was added to a final concentration of 1X to each sample and run on an SDS-PAGE gel, Coomassie stained, and imaged.

### PG purification and analysis

PG samples were analyzed as described previously (Alvarez et al., 2016; Desmarais et al., 2013). In brief, samples were boiled in SDS 5% for 2 h and sacculi were repeatedly washed with MilliQ water by ultracentrifugation (110,000 x g, 10 min, 20°C). The samples were treated with muramidase (100 μg/mL) for 16 hours at 37°C. Muramidase digestion was stopped by boiling and coagulated proteins were removed by centrifugation (10 min, 22,000 x g). The supernatants were first adjusted to pH 8.5-9.0 with sodium borate buffer and then sodium borohydride was added to a final concentration of 10 mg/mL. After reduction during 30 min at room temperature, the samples pH was adjusted to pH 3.5 with orthophosphoric acid. UPLC analyses of muropeptides were performed on a Waters UPLC system (Waters Corporation, USA) equipped with an ACQUITY UPLC BEH C18 Column, 130 Å, 1.7 μm, 2.1 mm X 150 mm (Waters, USA) and a dual wavelength absorbance detector. Elution of muropeptides was detected at 204 nm. Muropeptides were separated at 45°C using a linear gradient from buffer A (formic acid 0.1% in water) to buffer B (formic acid 0.1% in acetonitrile) in an 18-minute run, with a 0.25 mL/min flow.

To test the activity of EstG against cell wall substrates, sacculus or purified muropeptides were used as substrate. Reactions were performed in triplicates and contained 10 µg of purified enzyme, 50 mM Tris-HCl pH 7.5, 100 mM NaCl, and 10 µg of purified *Caulobacter* sacculus or 5 µg of purified M4, M5, D44 or D45, in a final 50 µL reaction volume. Reactions were incubated at 37°C for 24 h, then heat inactivated (100°C, 10 min) and centrifuged (22,000 x *g*, 15 min), for separation of soluble and pellet fractions. Soluble fractions were adjusted to pH 3.5. Pellet fractions were resuspended in water and further digested with muramidase for 16 h at 37°C. Muramidase reactions were reduced and adjusted to pH 3.5 as explained before. Both soluble and muramidase digested samples were run in the UPLC using the same PG analysis method described above.

Relative total PG amounts were calculated by comparison of the total intensities of the chromatograms (total area) from three biological replicas normalized to the same OD600 and extracted with the same volumes. Muropeptide identity was confirmed by MS/MS analysis, using a Xevo G2-XS QTof system (Waters Corporation, USA). Quantification of muropeptides was based on their relative abundances (relative area of the corresponding peak) normalized to their molar ratio. The program GraphPad PRISM® Software (Inc., San Diego, California, www.graphpad.com) was used for all statistical analyses. To determine the significance of the data, the t-test (unpaired) was performed.

### Crystallography, Data Collection, Structure Determination and Refinement

EstG protein purified for crystallography was prepared the same way as described above, with the exception of the storage buffer changed to 50 mM HEPES-NaOH pH 7.2, 150 mM NaCl, 1 mM DTT. Crystals of wild type EstG were grown by vapor diffusion in hanging drops set up with a Mosquito LCP robot (SPT Labtech, Melbourn, United Kingdom). Crystal growth was monitored using a crystallization imager ROCKIMAGER (Formulatrix, Bedford, Massachusetts). High quality crystals grew with a reservoir solution containing 20% PEG500 MME, 10% PEG20000, 0.1 M Tris/Bicine pH 8.5 and 90 mM mixture of sodium nitrate, sodium phosphate dibasic and ammonium sulfate (called EstG+SO_4_+TRS) or 20% PEG500 MME, 10% PEG20000, 0.1 M Tris/Bicine pH 8.5 and 100 mM mixture of DL-Alanine, Glycine, DL-Lysine and DL-Serine, (called EstG+TRS). Crystals grown in the first condition were soaked in 500 mM Tantalium bromide heavy metal solution for 1 hour (crystals called EstG + TaBr). Crystals were flash-cooled in mother liquor. Data of crystals of EstG +TRS (PDB ID 7UDA) were collected at National Synchrotron Light Source-II at beamline 17-ID-2 (FMX) on a Dectris EIGER X 16M while crystals of EstG in complex with SO_4_ and TRS (PDB ID 7UIC, EstG+SO_4_+TRS) and of EstG bound to tantalum bromide (PDB ID 7UIB, EstG+TaBr) were collected at 17-ID-1 (AMX) on a Dectris EIGER X 9M detector. Diffraction data were collected on a vector defined along the longest axis of the crystal (Miller et al., 2019). The datasets were indexed, integrated, and scaled using fastdp, XDS, and aimless (Kabsch, 2010). All EstG crystals belong to tetragonal space group and diffracted from 2.09 to 2.62 Å.

Since the N-terminal and C-terminal sequence of EstG differed from available homologous proteins, a model of EstG to use in molecular replacement was generated with the RoseTTAFold package (Baek et al., 2021). RoseTTAFold model weights as of July 16, 2021, UniRef30 clusters as of June 2020, PDB templates as of March 3, 2021, and the BFD (Steinegger and Söding, 2018) were used during model prediction. A C-terminal segment (Pro443-Arg462) that was predicted to extend as a random coil away from the molecular envelope was truncated from the model with the lowest predicted coordinate error to generate the final molecular replacement search model. The structure of EstG was determined by molecular replacement using PHASER (McCoy et al., 2007) with the RoseTTAFold model of EstG as a search model (Baek et al., 2021). The data were refined to a final resolution of 2.47, 2.09 and 2.62 Å using iterative rounds of refinement with REFMAC5 (Evans and Murshudov, 2013) and manual rebuilding in Coot (Emsley and Cowtan, 2004). Structures were validated using Coot (Emsley and Cowtan, 2004) and the PDB deposition tools. Each of the three models have more than 95 % of the residues in the preferred regions according to Ramachandran statistics (Table 2). Figures were render in PyMOL (v2.2.3, Schrödinger, LLC).

### Comparison with other beta lactamase binding proteins

A search using PDBeFOLD (Krissinel and Henrick, 2004) was conducted using EstG as a search model. Among them carboxyesterases, penicillin binding protein EstY29, and simvastatin synthase (PDBs 4IVK (Cha et al., 2013), 4P87 (Ngo et al., 2014), 3HLB (Gao et al., 2009)) aligned with root-mean-square deviations of 1.39, 1.62, 1.82 Å over 404, 387 and 400 amino acids, respectively. The structure of EstG was used to analyze and display the primary, secondary and quaternary structure of homologous proteins with ENDscript (Robert and Gouet, 2014).

### LCMS Analysis

All analysis was performed on a Dionex UHPLC and Q Exactive quadrupole Orbitrap system (Thermo Fisher, Waltham, Massachusetts). Two micrograms of each reaction and unreacted input was injected directly onto a HyperSil Gold C-18 2.1mm x 150mm reversed phase chromatography column. Analytes were separated using an increasing gradient that consisted of 0.1% formic acid in LCMS grade water as buffer A and 0.1% formic acid in LCMS grade acetonitrile as buffer B. Due to the hydrophilic nature of glucans, the gradient began with a 2-minute acquisition at 100% buffer A with a rapid ramp to 100% buffer B by 15 minutes before returning to baseline conditions for the remainder of the 20 minute experiment. The Q Exactive was operated in positive ionization mode using a data dependent acquisition method. An MS1 scan was acquired at 140,000 resolution with a scan range of 150 to 1500 m/z. The three most abundant ions from each MS1 scan were isolated for fragmentation using a three-step collision energy of 10, 30 and 100 and the fragment scans were obtained using 15,000 resolution. Ions with unassigned charge states or more than 3 charges were excluded from fragmentation. To prevent repeat fragmentation any ion within 5 ppm mass deviation of the selected ion was excluded from additional fragmentation for 30 seconds. The complete LCMS method in vendor .meth format and a text adaptation have been uploaded to LCMSMethods.org under the following DOI (dx.doi.org/10.17504/protocols.io.36wgq7djkvk5/v1). All Thermo .RAW instrument files have been uploaded to the MASSIVE public repository (Vizcaíno et al., 2014) under accession MSV000089142. The vendor .RAW files and processed results can be accessed during the review process using the following link: ftp://MSV000089142@massive.ucsd.eduand reviewer password EstG725.

### LCMS Data Analysis

All downstream data analysis was performed with Compound Discoverer 3.1 and Xcalibur QualBrowser 2.2 (Thermo Fisher). Briefly, all MS1 ions with a signal to noise of greater than 10:1 from the vendor .RAW files were considered for downstream analysis. The LCMS files were chromatographically aligned using an adaptive curve on all ions within a maximum mass shift of 2 minutes and with less than a 5 ppm mass discrepancy. The files were also normalized to compensate for concentration and loading differences between samples using a constant mean normalization. Ion identities were assigned using the mzCloud and ChemSpider databases using a maximum mass tolerance of 5ppm against library entries. In addition, a similarity search algorithm and custom compound class scoring module were used to flag ions that exhibited common glucose ions following fragmentation. Compounds of interest were flagged in the resulting output report by use of custom filter that eliminated ions that were of decreased abundance in the EstG reacted periplasm relative to both the unreacted periplast fraction and the periplast fraction treated with the EstG_S101A_.

### LCMS results

A total of 1,166 LCMS features were identified in the study. After removal of background signal and ions with an m/z of less than 600, 13 prospective ions were identified that appeared to be downregulated following incubation with the EstG protein. Of these molecules only one possessed a fragmentation pattern consistent with a glucan polymeric structure. This ion demonstrated an exact match by mass and an 83.7% fragment similarity to the cyclic hexasaccharide α-cyclodextrin. Figure 6C is a mirror plot that demonstrates the level of fragment sequence match between the fragmentation of this ion and α-cyclodextrin.

### Data availability

The final coordinates of EstG bound to TRS, EstG bound to SO_4_ and TRS, EstG bound to (Ta_6_Br_12_)^2^ have been deposited in the PDB with accession codes (7UDA, 7UIC and 7UIB) respectively.

## Acknowledgements

We thank the members of the Goley lab for helpful discussions and input. We thank Jean Marie Lacroix for helpful discussions about OPGs. We thank Patrick Viollier for SpmX antisera, Justine Collier for RelA’ plasmids, Martin Thanbichler for the periplasmic *blaM* plasmids, and Gyanu Lamichhane for providing nitrocefin. We thank Caren Freel Meyers, Natasha Zachara, Ronald Schnaar, Patrick Viollier, and Rico Rojas for helpful discussions regarding this work. We thank Patrick Viollier and Jordan Costafrolaz for initial discussions about CCNA_01638. We used Biorender.com to generate Figures 6 and 7. We would also like to thank BlaB, the original name of EstG, for being so fun to say for so many years.

This work is funded in part by the NIH, National Institute of General Medical Science through R35GM136221 (E.D.G.) R01GM108640 (E.D.G.), T32GM007445 (training grant support of A.K.D.). Mass spectrometry support from the Bumpus lab was supported in part by National Institutes of Health grant R01GM103853 (N.N.B.). Work at the AMX (17-ID-1) and FMX (17-ID- 2) beamlines is supported by the National Institutes of Health, National Institute of General Medical Sciences (P41GM111244), and by the DOE Office of Biological and Environmental Research (KP1605010), and the National Synchrotron Light Source II at Brookhaven National Laboratory is supported by the DOE Office of Basic Energy Sciences under contract number DE-SC0012704 (KC0401040). Research in the Cava lab is supported by The Swedish Research Council (VR), The Knut and Alice Wallenberg Foundation (KAW), The Laboratory of Molecular Infection Medicine Sweden (MIMS) and The Kempe Foundation. Research in the Chien lab is supported in part by the NIH, National Institute of General Medical Science through R35GM130320 (P.C.) and UMass NIH Chemistry Biology Interface Training Program T32GM008515 (R.Z.).

## Disclosures

S.B.G. is a founder and holds equity in AMS, LLC and is or was a consultant to Genesis Therapeutics, XinThera, and Scorpion Therapeutics.

---

**Figure 1--figure supplement 1:**
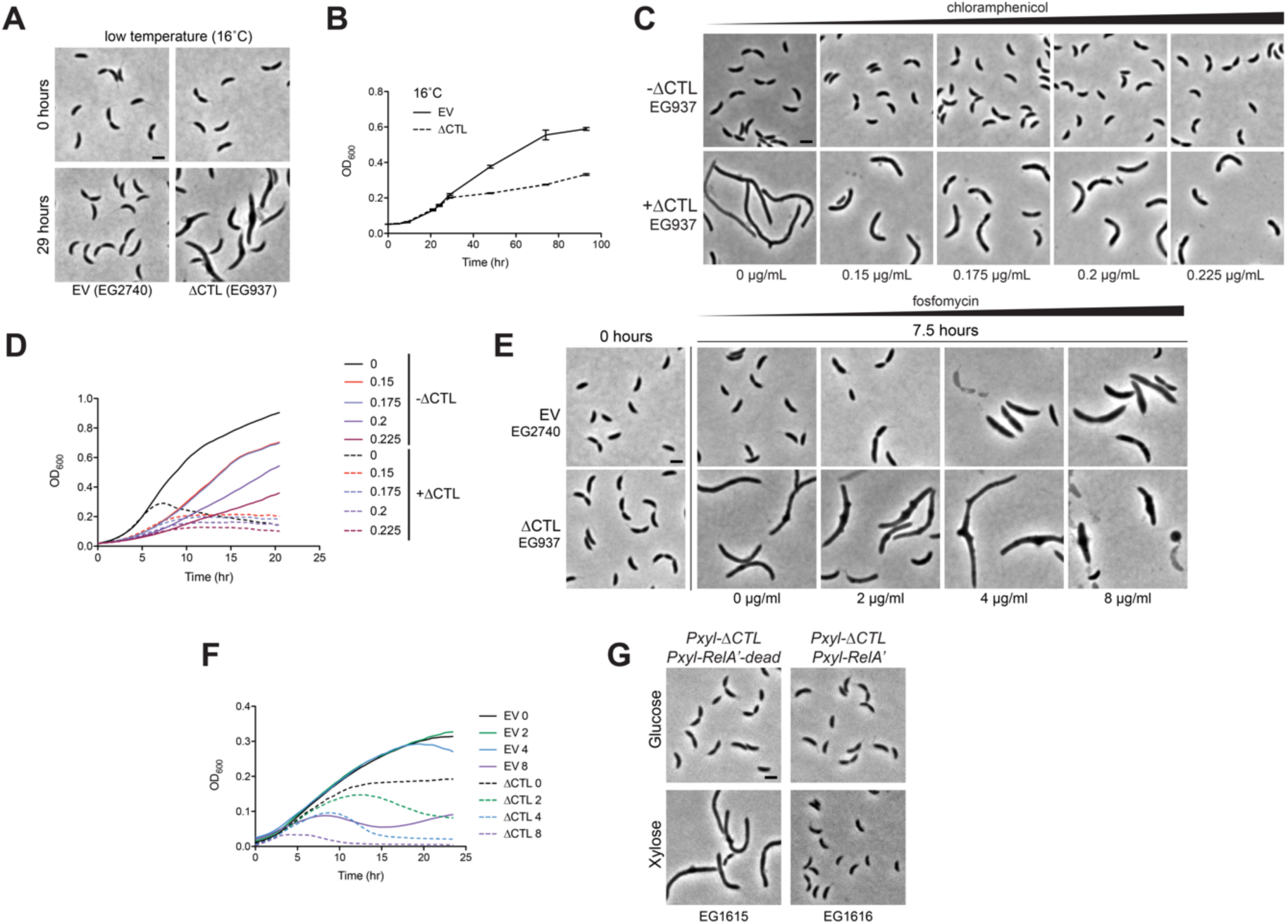
Slow growth does not suppress ΔCTL. **A.** Phase contrast images and **B.** growth curve of empty vector (EV, EG2740) and ΔCTL (EG937) grown with 0.3% xylose at 16°C to slow growth. **C.** Phase contrast images and **D.** growth curve of EG937 in the presence of 0.3% glucose (-ΔCTL) or 0.3% xylose (+ΔCTL) for 7.5 hours with increasing concentrations of chloramphenicol to slow growth. **E**. Phase contrast images and **F.** growth curve of EV and ΔCTL grown with 0.3% xylose with increasing concentrations of fosfomycin to slow growth. **F.** Phase contrast images of ΔCTL producing strains with xylose inducible RelA’ (high (p)ppGpp, EG1616) or catalytically dead RelA’dead (WT (p)ppGpp, EG1615) with 7 hours of 0.3% glucose or 0.3% xylose to induce ΔCTL and RelA’/RelA’dead. Scale bar, 2 µm.

**Figure 2--figure supplement 1:**
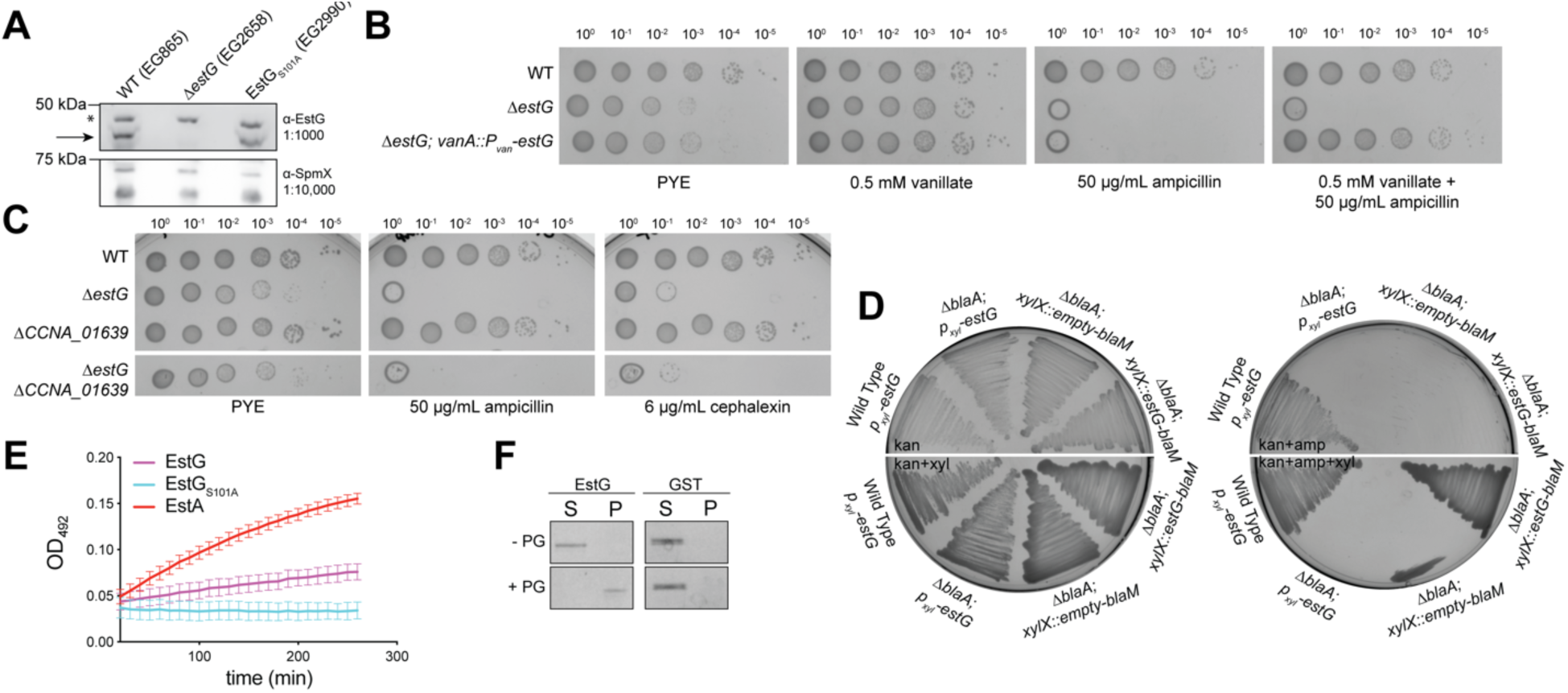
EstG is a periplasmic protein with broad antibiotic sensitivities. **A.** α-EstG (top) and α-SpmX (bottom) immunoblots of indicated strains at indicated dilutions. Arrow indicates band representing EstG. Asterisk denotes non-specific band. **B.** Spot dilutions of WT (EG865), Δ*estG* (EG2658), and Δ*estG* complemented with a vanillate inducible *estG* (EG3075) on PYE agar alone or with 0.5 mM vanillate, 50 µg/mL ampicillin, or both. Culture dilutions are as indicated. **C.** Spot dilutions of WT (EG865), Δ*estG* (EG2658), *ΔCCNA_01639* (EG3044), and *ΔestGΔCCNA_01639* (EG3047) on PYE agar alone or with 50 µg/mL ampicillin or 6 µg/mL cephalexin. Culture dilutions are as indicated. **D**. Periplasmic localization of EstG using indicated strains with fusions to *blaM* in a Δ*blaA* background. Cells are grown on PYE agar plates with indicated additives. Kanamycin 25 µg/mL (kan), 0.3% xylose, and ampicillin 50 µg/mL (amp). **E.** Nitrocefin hydrolysis of indicated proteins over time measured at OD_492_. **F.** Peptidoglycan (PG) binding ability of purified EstG or GST against wild type (WT) murein/sacculi. Upon ultracentrifugation, proteins unable to bind PG remain in the soluble fraction (S) and proteins that bind PG in the pellet (P).

**Figure 2--figure supplement 2:**
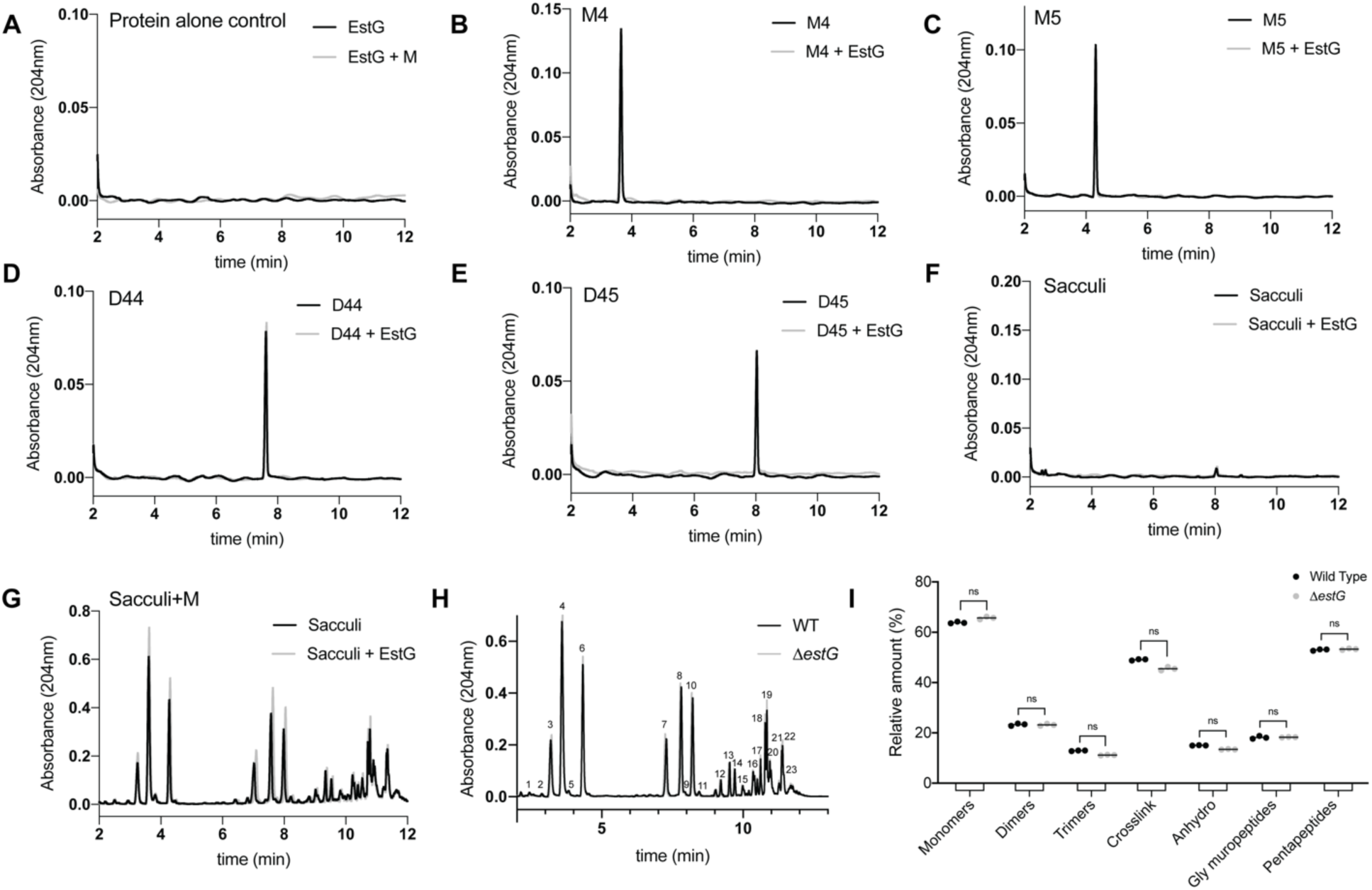
EstG does not have activity towards the peptidoglycan or its substituent moieties. *In vitro* reactions of EstG in the presence of **A.** protein alone, **B.** M4 (monomeric tetrapeptide), **C.** M5 (monomeric pentapeptide), **D.** D44 (dimeric tetrapeptide-tetrapeptide), **E.** D45 (dimeric tetrapeptide-pentapeptide), **F.** WT sacculi, and **G.** sacculi + muramidase treatment. **H.** Representative chromatograms of muropeptides prepared from WT (EG865) and Δ*estG* (EG2658). Relevant muropeptides are identified in Table 1. **I.** Relative molar abundance of the indicated muropeptide species from WT (EG865) and *ΔestG* (EG2658): monomers, dimers, trimers, crosslinkage, (1–6 anhydro) N-acetyl muramic acid containing muropeptides (anhydro, glycan chain termini), Gly containing muropeptides (Gly), and pentapeptides (penta).

**Figure 3--figure supplement 1:**
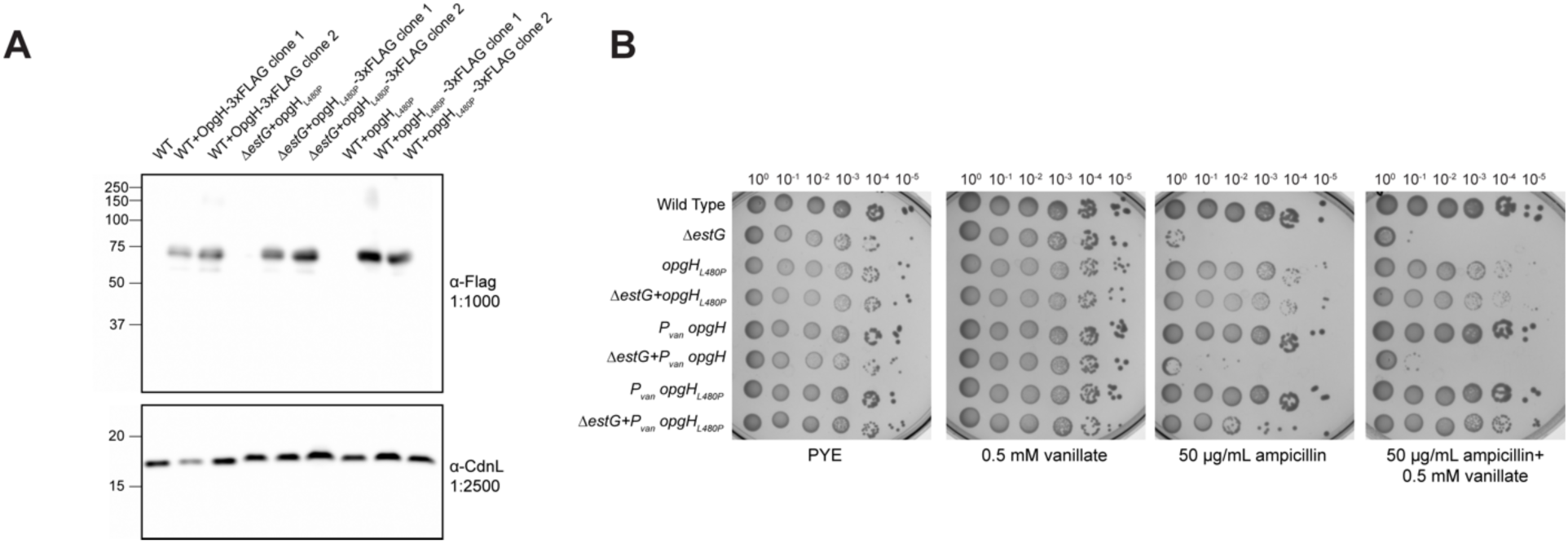
OpgH_L480P_ is not degraded and can suppress Δ*estG* sensitivity in a dominant fashion. **A.** α-Flag (top) and α-CdnL (bottom) immunoblots of indicated strains at indicated dilutions demonstrating the stability of a 3x-FLAG tagged variant of OpgH_L480P_. **B.** Spot dilutions on PYE agar alone or with added 0.5 mM vanillate and/or 50 µg/mL ampicillin of WT (EG865), *ΔestG* (EG2658), *opgH_L480P_* (EG3369), *ΔestG+opgH_L480P_* (EG3371), *P_van_-opgH* (EG3375), *ΔestG* + *P_van_- opgH* (EG3377), *P_van_-opgH_L480P_* (EG3440), *ΔestG+ P_van_-opgH_L480P_* (EG3442). Culture dilutions are as indicated.

**Figure 4--figure supplement 1:**
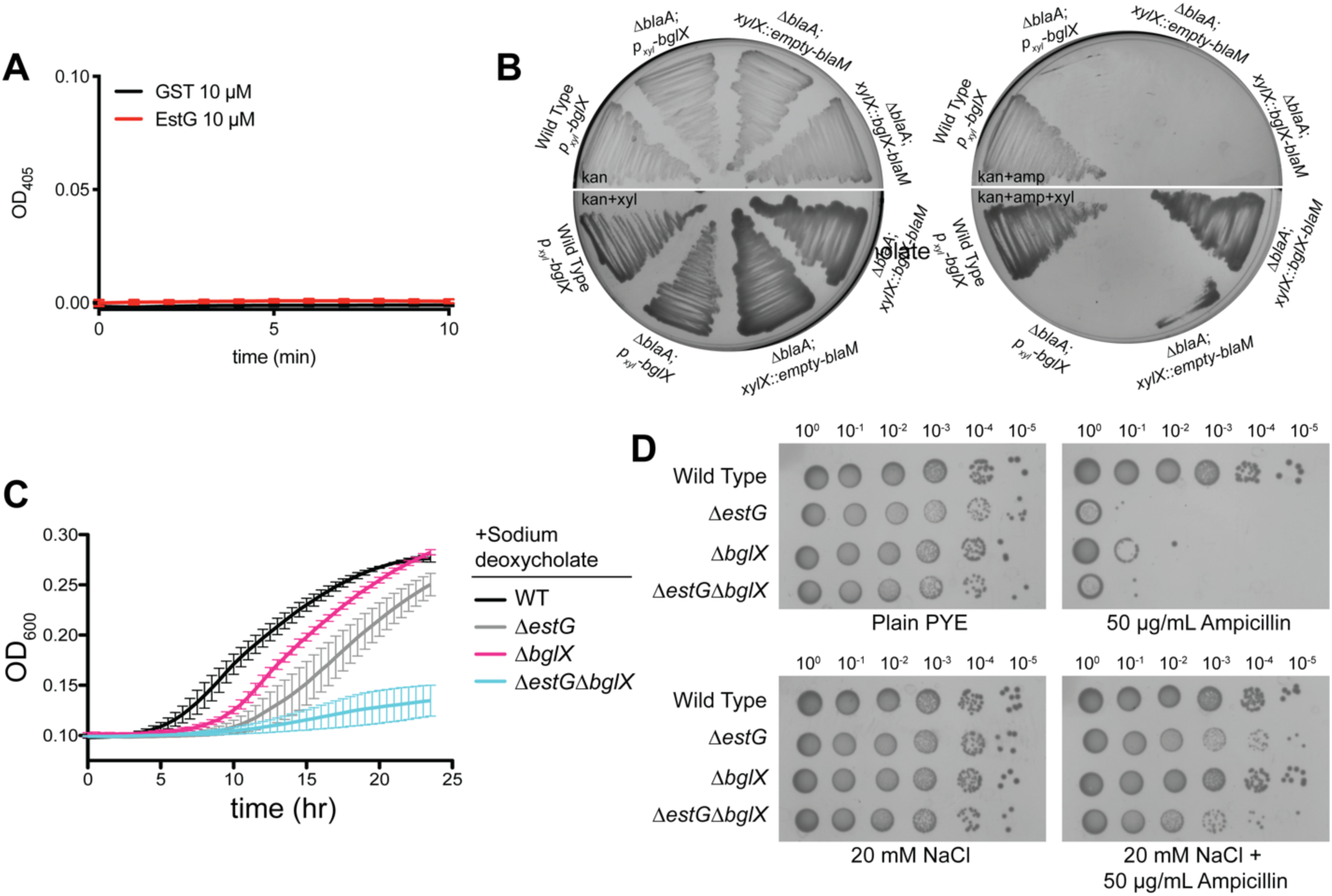
BglX localization and sensitivities are similar to EstG. **A.** 4-Nitrophenyl-β-D-glucopyranoside (pNPG) hydrolysis assay with 10 µM purified EstG or GST measured at OD_405_. **B.** Periplasmic localization of BglX using indicated strains with fusions to *blaM* in a Δ*blaA* background. Cells are grown on PYE agar plates with indicated additives. Kanamycin 25 µg/mL (kan), 0.3% xylose, ampicillin 50 µg/mL (amp). **C.** Growth curve of WT (EG865), Δ*estG* (EG2658), *ΔbglX* (EG3279), and *ΔestGΔbglX* (EG3282) with 0.6 mg/mL sodium deoxycholate. **D.** Spot dilutions of WT (EG865), Δ*estG* (EG2658), *ΔbglX* (EG3279), and *ΔestGΔbglX* (EG3282) on PYE agar alone or with added 50 µg/mL ampicillin and/or 20 mM NaCl. Culture dilutions are as indicated.

**Figure 5--figure supplement 1:**
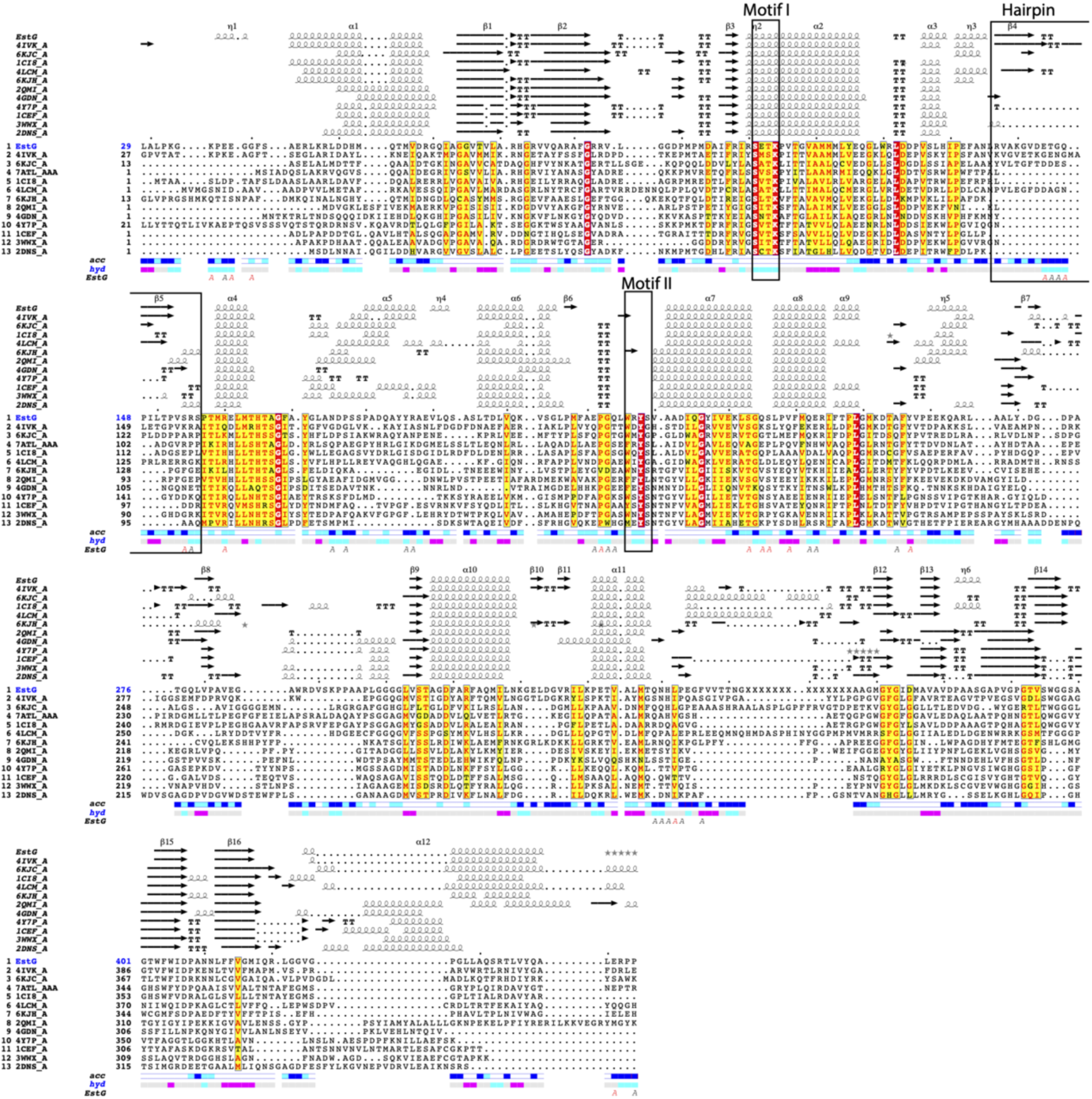
EstG has structural similarity to related enzymes. Multiple sequence and structural alignment of EstG and related enzymes displaying primary to quaternary structure information. The secondary structure elements are shown as helices, strands (arrows) and tight turns (TT). The sequence alignment is colored according to residue conservation with red background with white letter for identical, yellow background with red letters for conserved. Solvent accessibility (turquoise and yellow) and hydropathy scales per residue. Letter A indicates protein:protein interaction. The figure was done with ENDscript 2 (Robert and Gouet, 2014).

**Figure 5--figure supplement 2:**
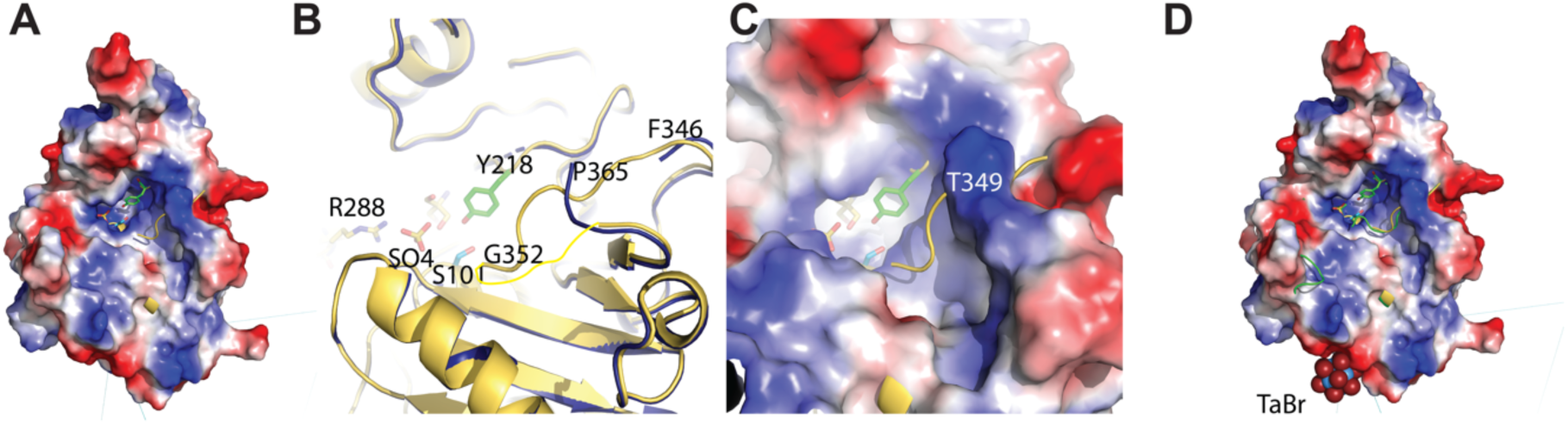
Loop present in EstG could be involved in catalysis. **A.** Electrostatic surface of EstG bound to Tris (TRS) structurally aligned to EstG bound to TRS and sulfate (SO_4_) displays the ordered loop F346 to G352 (yellow) and how it might occlude the binding site. Residues Ser101 and Tyr218 are shown in sticks. **B.** Zoom in and **C.** electrostatic surface rendering of the structural differences in loop 346-352. **D**. Structure of EstG bound to TRS structurally aligned to EstG+(Ta_6_Br_12_)^2^ (green cartoon). The tantalum bromide cluster, (Ta_6_Br_12_)^2^ is far from the loop, shown in spheres. EstG+(Ta_6_Br_12_)^2^ structure shows an alternative conformation for loop 272-276 and so protrudes from the surface.

**Figure 5--figure supplement 3:**
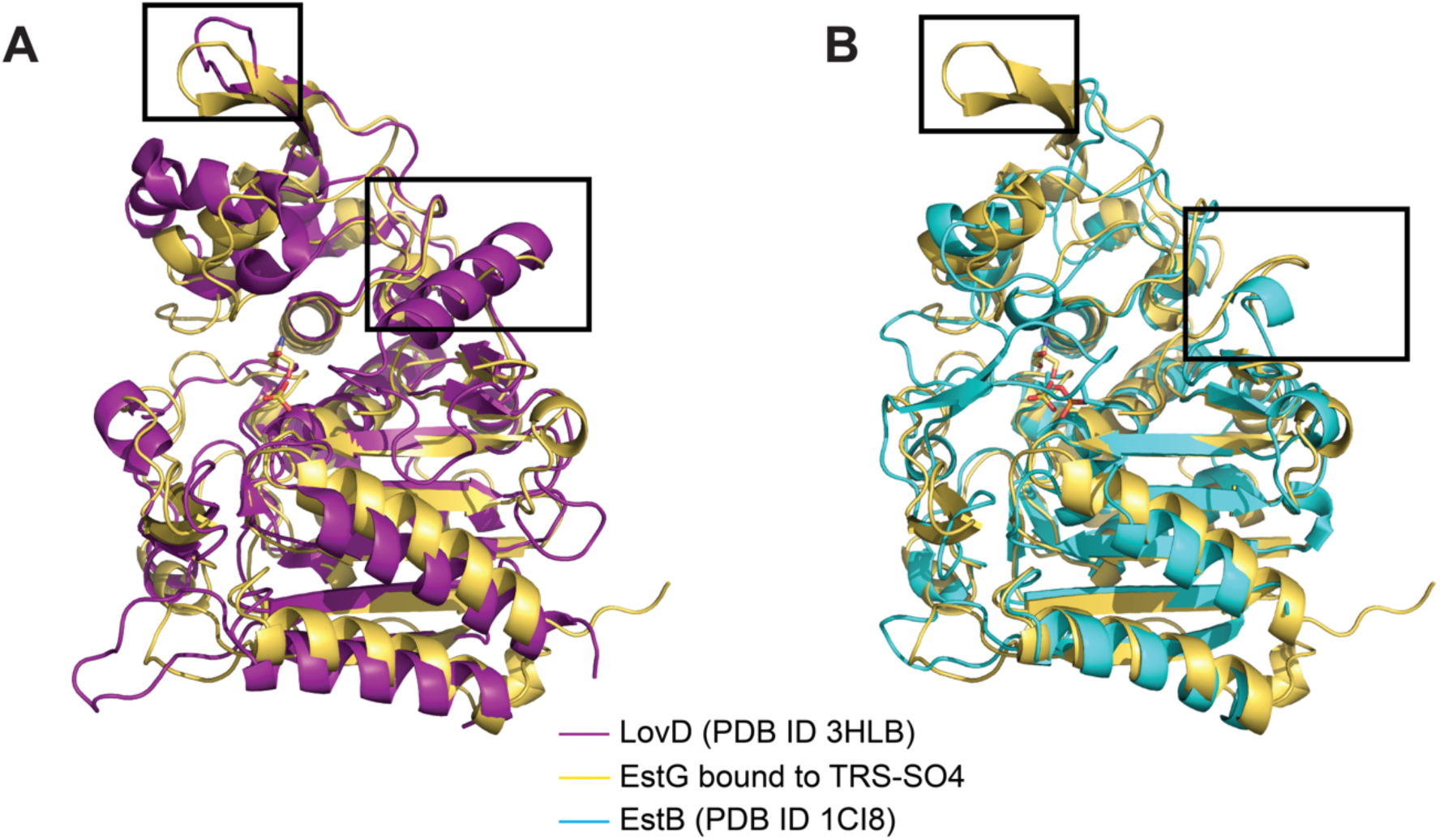
Structural alignment of EstG with related enzymes. **A.** Structural alignment of EstG with LovD (PDB ID 3HLB (Gao et al., 2009), purple) highlighting that EstG lacks the long helix between α11 and β12 (aa 340-350) present in 3HLB (aa 309-321). The hairpin insertion and the top of the hydrolase domain is in a different conformation. **B.** Structural alignment of EstG with EstB (PDB ID 1CI8 (Wagner et al., 2009), cyan) highlighting the insertion of the hairpin formed by β4 and β5 in EstG that is absent in EstB and in PDB IDs 4Y7P (Nakano et al., 2015), 1CEF (Kuzin et al., 1995), 3WWX (Arima et al., 2016), 2DNS (Okazaki et al., 2007).

**Figure 6--figure supplement 1:**
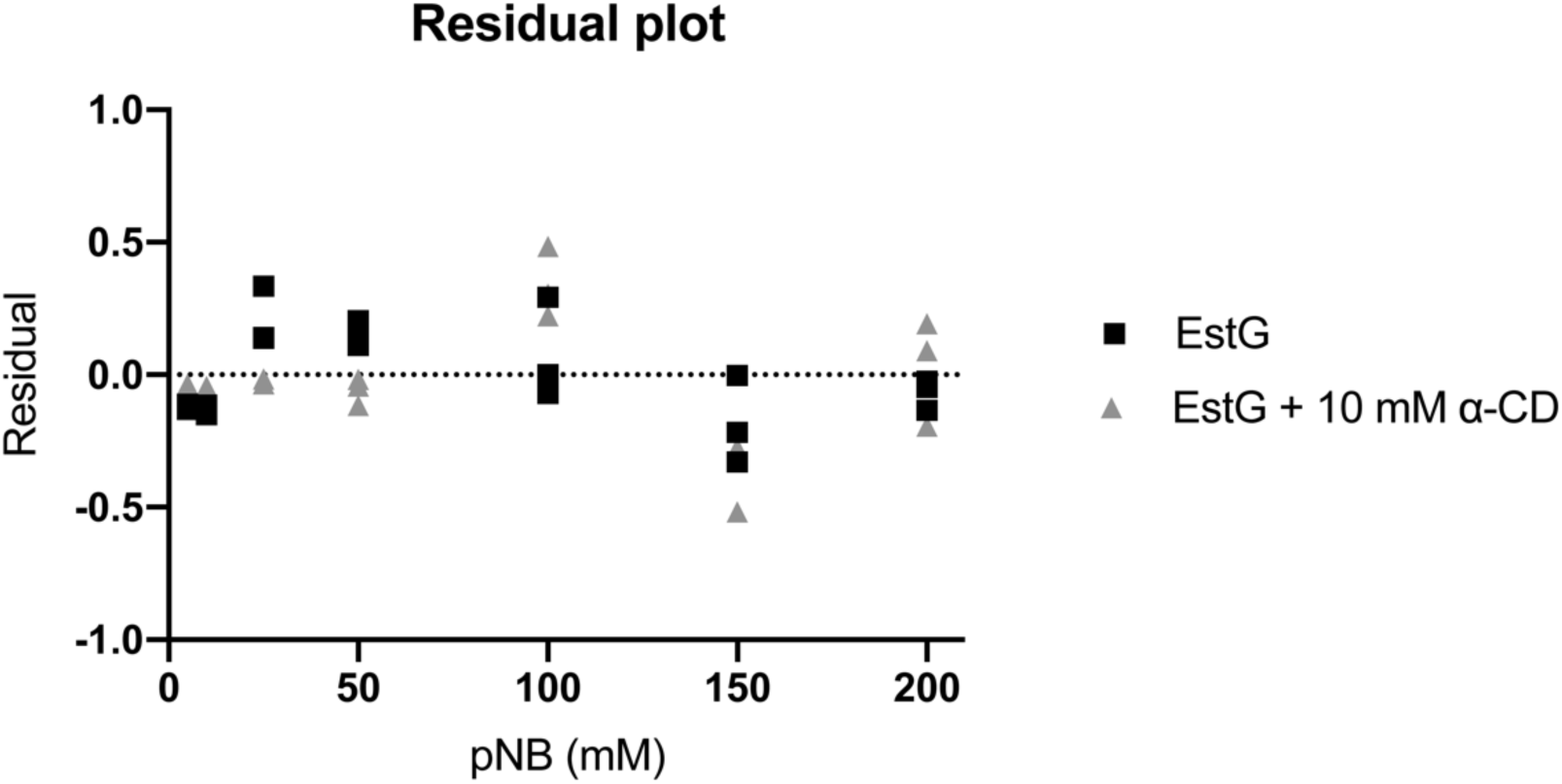
EstG residual in the presence and absence of α-cyclodextrin. Residual plot of the rate data from Figure 6E, presenting the deviation of the data points from the respective Michaelis-Menten fit.

**Supplement Table 1.**

Whole genome sequencing of suppressors for ΔCTL screen and Δ*estG* spontaneous suppressors.

**Supplement Table 2.**

Tn-Seq data for WT (EG865), RelA’ (EG1799), and ΔCTL+RelA’ (EG1616).

**Supplement Table 3.**

Strains and plasmids used in this study.

## Notes

### Competing Interest Statement

The authors have declared no competing interest.

